# Oncogene expression from extrachromosomal DNA is driven by copy number amplification and does not require spatial clustering

**DOI:** 10.1101/2022.01.29.478046

**Authors:** Karin Purshouse, Elias T. Friman, Shelagh Boyle, Pooran Singh Dewari, Vivien Grant, Alhafidz Hamdan, Gillian M. Morrison, Paul M Brennan, Sjoerd V. Beentjes, Steven M. Pollard, Wendy A. Bickmore

## Abstract

Extrachromosomal DNA (ecDNA) are frequently observed in human cancers and are responsible for high levels of oncogene expression. In glioblastoma (GBM), ecDNA copy number correlates with poor prognosis. It is hypothesized that their copy number, size and chromatin accessibility facilitate clustering of ecDNA and colocalization with transcriptional condensates, and that this underpins their elevated transcriptional activity. Here, we use super-resolution imaging and quantitative image analysis to evaluate GBM stem cells harboring distinct ecDNA species (*EGFR, MYC, PDGFR*). We found no evidence that ecDNA cluster with one another or closely interact with transcriptional condensates. Cells with *EGFR*-containing ecDNA have increased EGFR transcriptional output, but transcription per gene copy was similar in ecDNA compared to the endogenous chromosomal locus. These data suggest that is the increased copy number of oncogene-harbouring ecDNA that primarily drives high levels of oncogene transcription, rather than specific interactions of ecDNA with the cellular transcriptional machinery.

## Introduction

Glioblastoma (GBM) is characterized by intra-tumoral heterogeneity and stem cell-like properties that underpin treatment resistance and poor prognosis (Bulstrode et al., 2017; Suvà et al., 2014). GBM is divided into distinct transcriptional subtypes that span a continuum of stem cell/developmental and injury response/immune evasion cell states (Richards et al., 2021; Verhaak et al., 2010; Wang et al., 2021). Genetically, activation or amplification of *EGFR* (chr7) is altered in almost two-thirds of GBM (Brennan et al., 2013). Other commonly amplified genes include *PDGFRA* (chr4), *CDK4, MDM2* (chr12), *MET* and *CDK6* (chr7) with multicopy extra-chromosomal DNA (ecDNA) considered a major mechanism for oncogene amplification (Brennan et al., 2013; Kim et al., 2020).

EcDNA are a particularly prominent feature of GBM, with 90% of patient-derived GBM tumor models harboring ecDNA (Turner et al., 2017). However, there is much broader interest in mechanisms of ecDNA function across many solid tumours, as ecDNA enable rapid oncogene amplification in response to selective pressures, and have been shown to correlate with poor prognosis and treatment resistance (Kim et al., 2020; Nathanson et al., 2014; Vicario et al., 2015). EcDNA are centromere-free DNA circles of around 1-3Mb in size that frequently exist as doublets (double minutes), but also as single elements (Verhaak et al., 2019). Although ecDNA were previously identified in 1.4% of cancers, more recent studies have shown their prevalence to be significantly higher (Fan et al., 2011; Kim et al., 2019; Turner et al., 2017). EcDNA can lead to oncogene copy number being amplified to >100 in any given cell, with significant copy number heterogeneity between cells (Lange et al., 2021; Turner et al., 2017). Freed from the constraints imposed by being embedded within a chromosome, ecDNA have spatial freedom and can adapt to targeted therapeutics (Lange et al., 2021; Nathanson et al., 2014). For example, the EGFR variant *EGFRvIII* (exon 2-7 deletion) is found on ecDNA, and is associated with an aggressive disease course and resistance mechanisms against EGFR inhibitors (Brennan et al., 2013; Inda et al., 2010; Nathanson et al., 2014; Turner et al., 2017).

As well as their resident oncogenes, ecDNA also harbor regulatory elements (enhancers) required to drive oncogene expression (Morton et al., 2019; Zhu et al., 2021). Consistent with this, ecDNA have been found to have regions of largely accessible chromatin (assayed by ATAC-seq), indicative of nucleosome displacement by bound transcription factors, and to be decorated with histone modifications associated with active chromatin (Wu et al., 2019). Transcription factors densely co-bound at enhancers have been suggested to nucleate condensates or ‘hubs’ (Cho et al., 2018; Rai et al., 2018; Strom and Brangwynne, 2019), enriched with key transcriptional components such as Mediator and RNA polymerase II (PolII) to drive high levels of gene expression (Cho et al., 2018; Chong et al., 2018; Sabari et al., 2018). Given the colocation of enhancers and driver oncogenes on ecDNA, it has therefore been suggested that ecDNA may cluster together in the nucleus, driving the recruitment of a high concentration of RNA PolII and creating ecDNA-driven nuclear condensates that in turn enhance the transcriptional output from ecDNA (Adelman and Martin, 2021; Hung et al., 2021; Yi et al., 2021; Zhu et al., 2021).

Here, using super-resolution imaging in primary GBM cell lines, we find that ecDNA are widely dispersed throughout the nucleus and we find neither evidence of ecDNA clustering together, nor any significant spatial overlap between ecDNA and PolII condensates. As expected, we show that expression from genes on ecDNA, both at mRNA and protein level, correlates with ecDNA copy number in the tumor cell lines. However, transcription of oncogenes present on each individual ecDNA molecule appears to occur at a similar efficiency (transcripts per copy number) to that of the equivalent endogenous chromosomally located gene. These data suggest it is primarily the increased copy number of ecDNA, and not a specific property of nuclear colocalization, that drives the increased transcriptional capacity of their resident oncogenes.

## Results

### EcDNA are more frequently located centrally in the nucleus in GBM cells

We characterized two GBM-derived Glioma Stem Cell (GSC) primary cell lines containing multiple *EGFR*-harboring ecDNA (ecEGFR) populations (GCGR-E26 and GCGR-E28, referred to here as E26 and E28). Whole genome sequencing (WGS) analysis using Amplicon Architect (Deshpande et al., 2019) indicated that E26 ecDNA harbor an *EGFRvIII* (exon 2-7 deletion), and E28 have a subpopulation of ecDNA with *EGFR* exon 7-14 deleted (Figure S1A). The presence of *EGFR* on ecDNA was confirmed by DNA FISH on metaphase chromosomes (Figures 1A and 1B). E26 harbored more ecDNA per cell than E28 (Figure 1C), with approximately 10% of metaphases also indicating the presence of a chromosomal homogeneously staining region (HSR) (Figure 1A; arrow).

**Figure 1.**
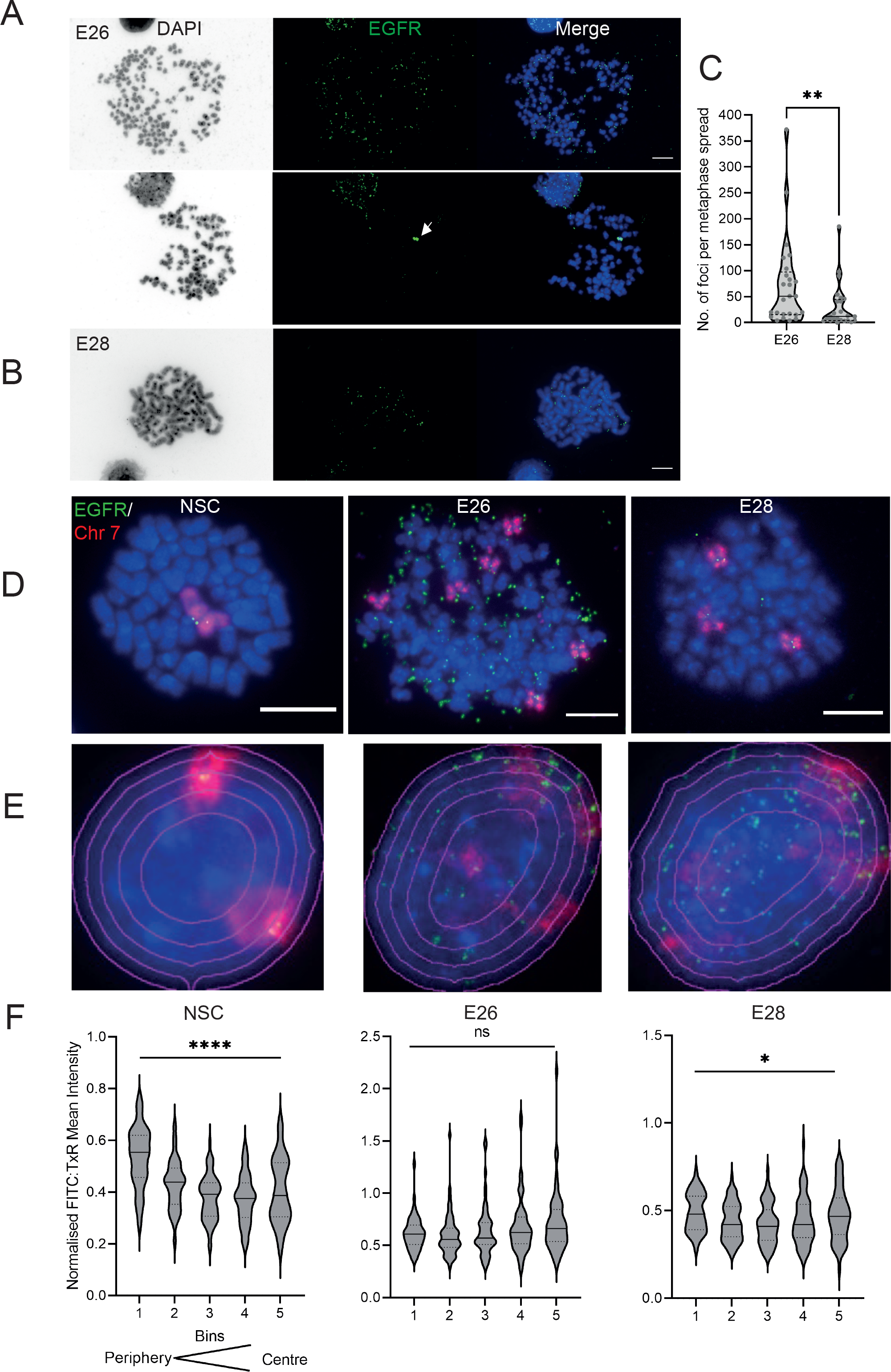
The nuclear localization of ecDNA in GBM cell lines. A) DNA FISH on metaphase spread of the E26 cell line showing EGFR (green) present on ecDNA, and on an HSR (arrowed) detected in 10% of metaphases. Scale bar = 10 μm. B) As for (A) but for the E28 cell line. C) Violin plot of the number of EGFR DNA FISH signals per metaphase spread of E26 and E28 cells. Median and quartiles are shown. ** p<0.01. D and E) Representative DNA FISH images of metaphase spread (D) and 2D nuclei (E) for NSC, E26 and E28 cells showing signals for chromosome 7 (red) and EGFR (green). Scale bar = 10 μm. The 5 erosions bins from the periphery to the centre of the3 nucleus are shown in E. F) EGFR FISH signal intensity normalised to that for chromosome 7 (FITC:TxR Mean Intensity) across 5 bins of equal area eroded from the peripheral (Bin 1) to the centre (Bin 5) of the nucleus for NSC, E26 and E28 cell lines. >55 nuclei per cell line. Median and quartiles shown. ns = not significant, * p<0.05, FITC and TxR signal normalised to DAPI shown in Supplementary Figure S1A. Statistical data relevant for this figure are in Table S1.

Human chromosomes have non-random nuclear organization, with active regions preferentially located toward the central regions of the nucleus (Boyle et al., 2001; Croft et al., 1999). We sought to determine the nuclear localization of ecDNA in GBM cell lines as compared with the endogenous chromosomal *EGFR*. Endogenous *EGFR* is located on human chromosome 7, a chromosome generally found toward the periphery of the nucleus (Boyle et al., 2001). DNA FISH for chromosome 7 and *EGFR* in nuclei from human foetal neural stem cells (NSCs) confirmed this (Figures 1D, 1E, S1B and S1C). Metaphase spreads of the two tumor lines showed 3-6 copies of chromosome 7 in E26 and frequently 3 copies in E28 (Figure 1D). In interphase nuclei (Figure 1E), signal intensity analysis for equally sized bins eroded from the edge to the center of each nucleus indicated chromosome 7 and *EGFR* signal intensity was preferentially located toward the nuclear periphery in each cell lines (Figure S1B and S1C). Even once chromosome 7 signal was accounted for, *EGFR* DNA FISH signal was still highest in the periphery of NSC nuclei and lowest in the central regions (p<0.0001) (Figure 1F), likely reflecting the centromere proximal localization of endogenous *EGFR* on chromosome 7 (Carvalho et al., 2001). This radial organization was still significant (p=0.0117), but much less marked, in E28 cells which have on average a modest number of EGFR ecDNA compared to endogenous copies (Figure 1C). In E26 cells, which have a very high copy number of ecDNA, this preference for a more peripheral localization is lost (p=0.0598). These data suggest that, freed of the constraints on nuclear localization imposed by human chromosome 7, *EGFR* genes located on ecDNA can access more central regions of the nucleus.

### EcDNA do not cluster in the nucleus

It has been suggested that ecDNA cluster into “ecDNA hubs,” within nuclei of cancer cells, including for *EGFRvIII*-containing ecDNA in other GBM cell lines (HK359 and GBM39) (Hung et al., 2021; Yi et al., 2021). We sought to quantify this using our E26 and E28 GBM cells with a single oncogene-harboring ecDNA population (*EGFR* variant amplicons) (Figure 2A). We used 3D image-based analysis of the *EGFR* DNA FISH signals to determine if there is clustering of ecDNA. Despite the difference in ecDNA copy number between the two GBM cell lines (Figure 1C), the mean shortest interprobe distance per nucleus was routinely >1.5μm in both cell lines (Figure 2B). The single shortest interprobe distance per nucleus was also larger (0.24μm, E26; 0.25μm, E28) than would be expected if there were clustering of ecDNA in the close proximity required for coordinated transcription in condensates or hubs; this should be ∼200nm or less (Figure 2C).

**Figure 2.**
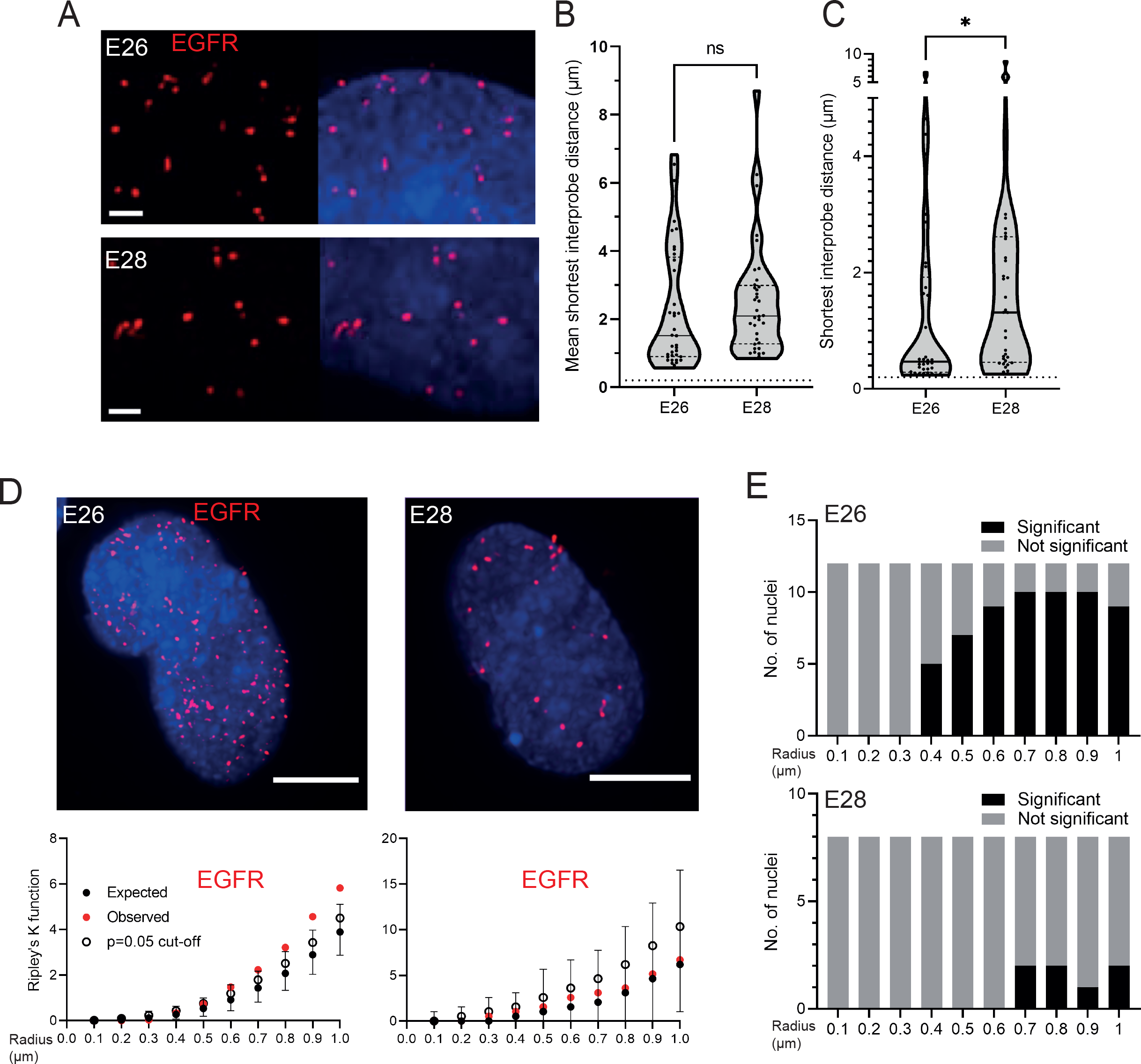
EGFR-containing ecDNA do not cluster in the nucleus. A) Representative images shown as maximum intensity projection of DNA FISH for EGFR (red) in the nuclei of E26 (top) and E28 (bottom) GBM cell lines, scale bar = 1 μm. B) Violin plots showing mean shortest interprobe distance between EGFR foci per nucleus in E26 and E28 cell lines. Dotted line denotes y=200nm. Number of nuclei (n): E26 = 37, E28 = 36. C) As for (B) but for shortest single distance between two EGFR foci in any nucleus. n: E26 = 37, E28 = 36. Statistical significance examined by Mann-Whitney test. ns = not significant, * p<0.05 and are detailed in Table S2. D) (top) Representative maximum intensity projection images of EGFR DNA FISH (red) in nuclei of E26 and E28 cells. Scale bar = 5 μm. (bottom) Associated 3D Ripley’s K function for these nuclei showing observed K function (red), max/min/median (black) of 10,000 null samples with p=0.05 significance cut-off shown (empty black circle). E) Ripley’s K function for EGFR DNA FISH signals showing number of E26 and E28 nuclei with significant and non-significant clustering at each given radius. p values were calculated using Neyman-Pearson lemma with optimistic estimate p value where required (see Methods), and Benjamini-Hochberg Procedure (BHP, FDR = 0.05).

The analysis above quantified distances between spots but does not allow for determination of whether there is a non-random distribution of foci in the nuclei. We therefore used 3D Ripley’s K function to determine the observed spatial pattern of the foci in each nucleus and compared this with a random null distribution of 10,000 simulations of the same number of foci in the same volume. We powered this to identify any significant clustering at each radius in 0.1μm increments between 0.1-1μm (examples of E26 and E28 nuclei and their corresponding Ripley’s K function in Figure 2D). The E26 cell line had some nuclei with significant non-random distribution of ecDNA, but only at >400nm radial distances, and E28 only had occasional nuclei with significant non-random distribution of ecDNA at >700nm (Figure 2E). These data therefore do not suggest any clustering at distances that might suggest colocalization of *EGFR* ecDNA, and hence coordinated transcription.

To ensure this was not because ecDNA were so tightly clustered that they could not be resolved by FISH, we analyzed another primary GBM cell line (termed E25) which has two different oncogenes carried on separate ecDNA populations: *CDK4* and *PDGFRA (*Figures S2A and B). E25 cells were also found to have no obvious clustering of the two ecDNA populations (Figure 3A). Indeed, the mean shortest inter-probe distances per nucleus were overwhelmingly >1 μm, suggesting ecDNA were generally not in close proximity (Figure 3B). The shortest interprobe distances for *CDK4-CDK4* and *CDK4-PDGFRA* were shorter than for *PDGFRA-PDGFRA* foci, as expected given the higher copy number of *CDK4* ecDNA (Figure S2B); however, there was no significant difference in the shortest distance between *CDK4-CDK4* and *PDGFRA-CDK4* foci (Figure 3C). No two *CDK4* or two *PDGFRA* foci were <200nm apart, and only 4 *CDK4-PDGFRA* distances were <200nm (4/1011 (0.39%) *CDK4* foci, 4/518 (0.77%) *PDGFRA* foci) (Figure 3C). These data suggest that clustering is not a significant feature of two separate populations of ecDNA.

**Figure 3.**
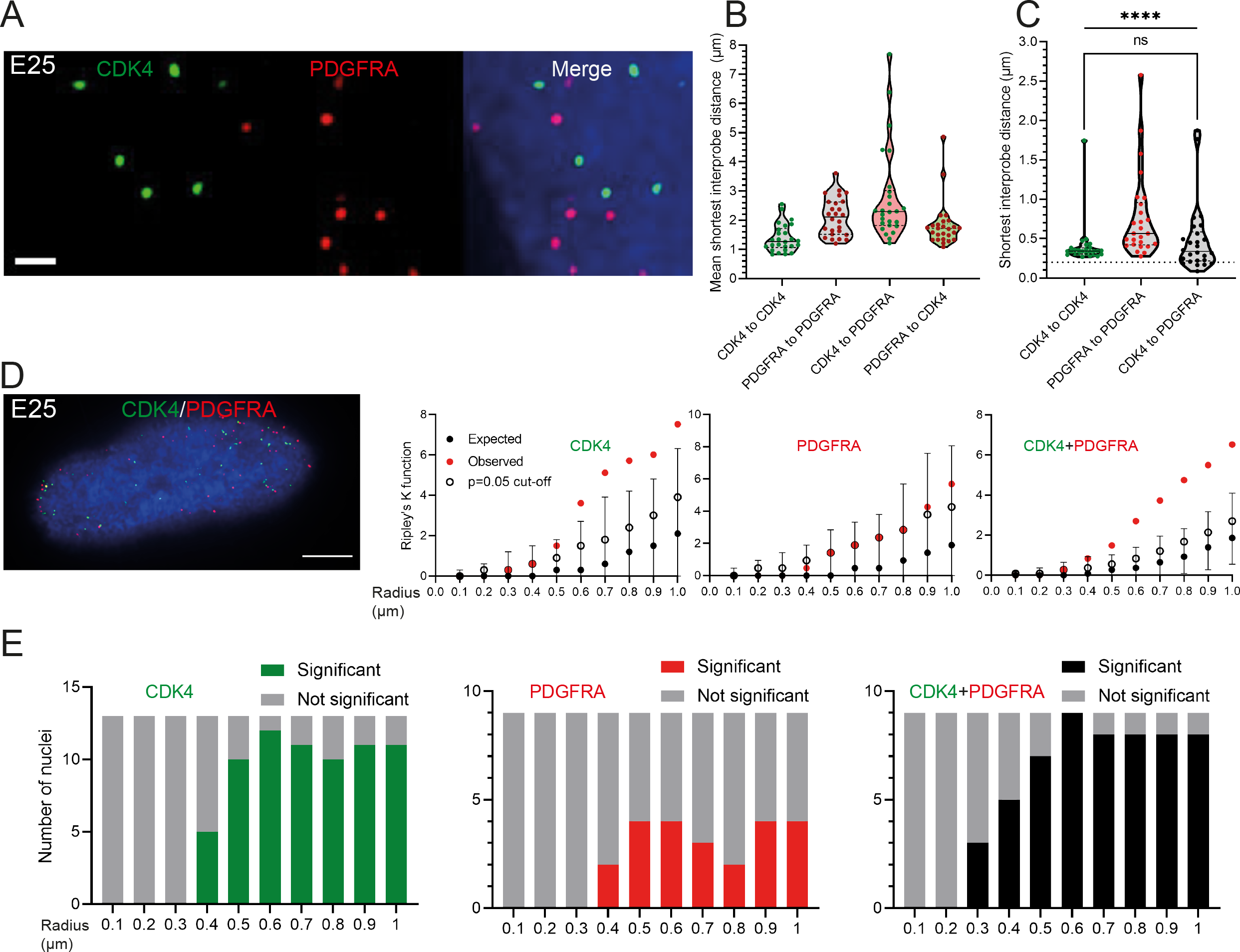
Two separate ecDNA populations do not cluster in the nucleus. A) Representative maximum intensity projection images of DNA FISH for CDK4 (green) and PDGFRA (red) in a E25 nucleus. Scale bar = 1 μm. B) Violin plots showing mean shortest distance between CDK4 and PDGFRA foci per E25 nucleus. Dotted line denotes y=200nm. C) As for (B) but showing the shortest single inter-probe distance measured in any nucleus, Statistical significance examined by Mann-Whitney Test (hooked line, ns = not significant) and Kruskall-Wallis (straight line, **** p<0.0001). Statistical data are detailed in Table S3. D) (left) Representative maximum intensity projection image shown of E25 nucleus hybridized with probes for CDK4 (green) and PDGFRA (red). Scale bar = 5 μm. (right) Ripley’s K function for this nucleus showing observed K function (red), max/min/median (black) of 10,000 null samples with p=0.05 significance cut-off shown (empty black circle) for CDK4, PDGFRA, and CDK4 and PDGFRA spots combined. E) Ripley’s K function for E25 nuclei showing number of nuclei with significant and non-significant clustering at each given radius for CDK4 spots, PDGFRA spots and CDK4 and PDGFRA spots combined. p values were calculated using Neyman-Pearson lemma with optimistic estimate p value where required (see Methods), and Benjamini-Hochberg Procedure (BHP, FDR = 0.05).

We used 3D Ripley’s K function to evaluate point patterns in this dual ecDNA oncogene E25 cell line (Figure 3D). Some nuclei had significant non-random distribution of *PDGFRA* ecDNA at >400nm, and most nuclei had non-random distribution of *CDK4* ecDNA at >400nm (Figure 3E). When both foci were combined, there was no significant clustering at <300nm in any nucleus, and the number of nuclei with significant non-random distribution at a given radius rose with increasing radial distance (Figure 3E). The analysis was repeated with a smaller (150nm diameter) spot size to ensure no small FISH foci were omitted that might skew our analysis. There were no instances where clustering occurred in any of the GBM cell lines or spot combinations at <300nm (Figures S3A and 3B).

DNA FISH detects all ecDNA, so it might be that only transcriptionally active elements cluster. Therefore, we repeated the clustering analysis using RNA FISH to detect nascent *EGFR* transcripts in the nuclei of E26 cells. As for DNA FISH, we detected no evidence of clustering at <400nm (Figures S3C and S3D). Overall, these data suggest that ecDNA do not colocalize in the nucleus more than expected by chance.

### Transcriptional condensates are few and do not colocalize with ecDNA

We next assessed whether ecDNA foci colocalize with high local concentrations of the transcriptional machinery – transcription condensates – to create ecDNA transcription hubs (Figure 4A). First, we examined the presence of condensates by immunofluorescence for RPB1 (POLR2A), the largest subunit of RNA PolII. We found that condensates were sparse with only a few clearly visible per nucleus and some nuclei harboring no condensates (Figures 4B and 4C).

**Figure 4.**
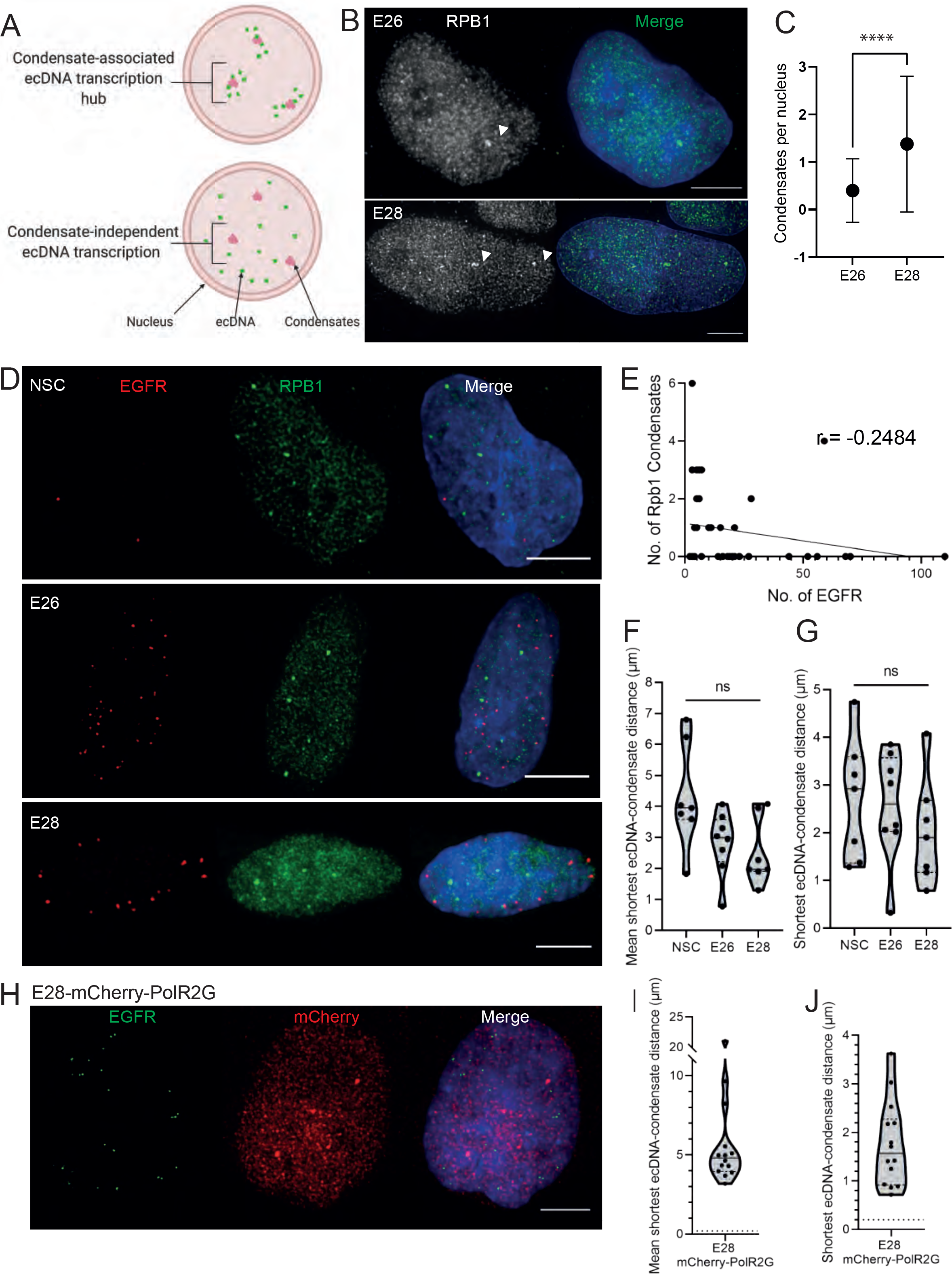
ecDNA do not colocalize with large condensates of the transcriptional machinery. A) Cartoon to illustrate condensate-associated ecDNA transcription hubs and condensate-free ecDNA transcription. Created with BioRender.com. B) Representative images of RNA polymerase II (RPB1) condensates (arrow heads) detected by immunofluorescence. Scale bar = 5 μm. C) Number of RPB1 condensates (mean +-SD) detected per nucleus in E26 and E28 cell lines. **** p <0.0001. D) Representative maximum intensity projection images of immunoFISH in NSC, E26 and E28 cell lines: Immunofluorescence for Rpb1 (green) and EGFR DNA FISH (red). Scale bar = 5 μm. E) Spearman’s correlation between number of EGFR ecDNA foci and number of Rpb1 condensates, p = 0.13. F) Violin plot of mean shortest interprobe distance per nucleus between EGFR foci and PolII condensates in NSC, E26 and E28 cell lines. G) As for (F) but for shortest single distance in each nucleus. ns, not significant. H) Representative images of immunoFISH in the E28 mCherry-Pol2RG cell line: Immunofluorescence for mCherry and EGFR DNA FISH. Scale bar = 5 μm. I) Violin plot of mean shortest distance per nucleus between EGFR foci and condensates detected by Pol2RG-mCherry fusion. Dotted line denotes y=200nm. J) As for (I) but for shortest single distance in each nucleus. Statistical data relevant for this figure are in Table S4.

We used 3D analysis of immunoFISH in NSCs and compared these to E26 and E28 GBM cells to establish whether ecDNA and condensates colocalized (Figure 4D). We found no correlation between the number of condensates and the number of ecDNA (Figure 4E). The mean shortest distance between EGFR foci and condensates per nucleus was routinely > 1 μm in all cell lines, despite the greater number of *EGFR* foci in the GBM cell lines (Figure 4F). The single shortest distance per nucleus between an *EGFR* locus and a condensate was not significantly different across NSC and tumor lines (Figure 4G), suggesting no clustering of ecDNA at condensates. There were no instances where the distance between ecDNA and condensates was <200nm. To test if this was also the case for the nascent *EGFR* RNA transcript, we repeated this analysis using nascent RNA FISH, and observed the same result (Figure S4A-C). As the distance distributions to RPB1 foci were similar for DNA and RNA FISH, this suggests that close proximity to RPB1 condensates does not alter the probability that the ecDNA is transcribed.

To ensure this result was not specific to this PolII antibody, we repeated this analysis using E28 cells in which mCherry had been fused to endogenous PolR2G, a key subunit of RNA PolII (Cramer et al., 2000), by knock-in (Figure 4H). The mean distance between ecDNA foci and condensates (Figure 4I) and the shortest minimum distance in any given nucleus (Figure 4J) further support that there is no close spatial relationship apparent between ecDNA and transcriptional condensates.

### Levels of transcription from ecDNA reflect copy number but not enhanced transcriptional efficiency

Having shown a lack of colocalization of ecDNA, either with each other or with PolII condensates, we proceeded to characterize the levels of expression of *EGFR* on ecDNA. Flow cytometry using the EGFR ligand conjugated to a fluorophore (EGF-647) revealed consistently higher levels of EGFR in the GBM cells than NSC, with highest signal in E26 (Figures S5A and S5B), consistent with the higher ecDNA copy number in E26 GBM cells than in E28 (Figure 1C). To confirm this link between ecDNA number and levels of EGFR, E26 and E28 cells were sorted by FACS into EGFR-high and EGFR-low populations (Figure S5C). In both tumor cell lines, DNA FISH demonstrated that EGFR-high cells had a significantly higher number of *EGFR* DNA foci than EGFR-low (Figures S5D and S5E). We next sought to characterize the transcriptional efficiency (per copy number) of chromosomal and ecDNA-located *EGFR* genes by nascent RNA FISH using a probe targeting the first intron of *EGFR*. As expected, nascent RNA FISH foci were more frequent in the *EGFR* ecDNA-harboring cell lines than in NSCs and were more frequent in the E26 GBM cell line than in E28 (Figures S5F and S5G).

Previous work has shown that ecDNA chromatin is highly accessible with greater transcript production per oncogene than chromosomal loci (Wu et al., 2019). We hypothesized that combining nascent RNA FISH and DNA FISH would reveal evidence of greater transcriptional efficiency with a higher RNA:DNA *EGFR* FISH foci ratio in cells with higher ecDNA copy number. We performed nascent *EGFR* RNA and *EGFR* DNA FISH to test this hypothesis (Figure 5A). When comparing the RNA:DNA ratio of all nuclei, surprisingly, there was no significant difference between NSC and E28, and both had a significantly lower ratio than E26 (Figure 5B). This suggests differences in transcriptional efficiency between the two tumor cell lines, although both showed an apparently linear correlation between the number of nascent *EGFR* RNA and *EGFR* DNA foci per nucleus (Figure 5C). To explore whether any differences in *EGFR* transcription in these cell lines could be due to ec*EGFR-*driven increased transcriptional efficiency, we sought to separate nuclei into those with predominantly chromosomal and extrachromosomal EGFR based on chromosome 7 number in these cell lines (Figure 1D). We categorized nuclei that had <10 *EGFR* DNA foci as being predominantly chromosomal *EGFR*-harboring, and >10 *EGFR* foci as being predominantly ecDNA EGFR-harboring, and compared the RNA:DNA foci number ratio. Unexpectedly, these were not significantly different (Figure 5D). This was repeated for E28 using <6/>6 foci as the cut-offs (to allow for the lower chromosome 7 copy number in this cell line), and this also showed no significant difference (Figure S5H). This suggests that within the same cell population, EGFR transcription from ecDNA and chromosomes occurs at similar levels when normalized to copy number. There is no increased transcriptional efficiency from ecDNA than from chromosomal DNA based on these analyses.

**Figure 5.**
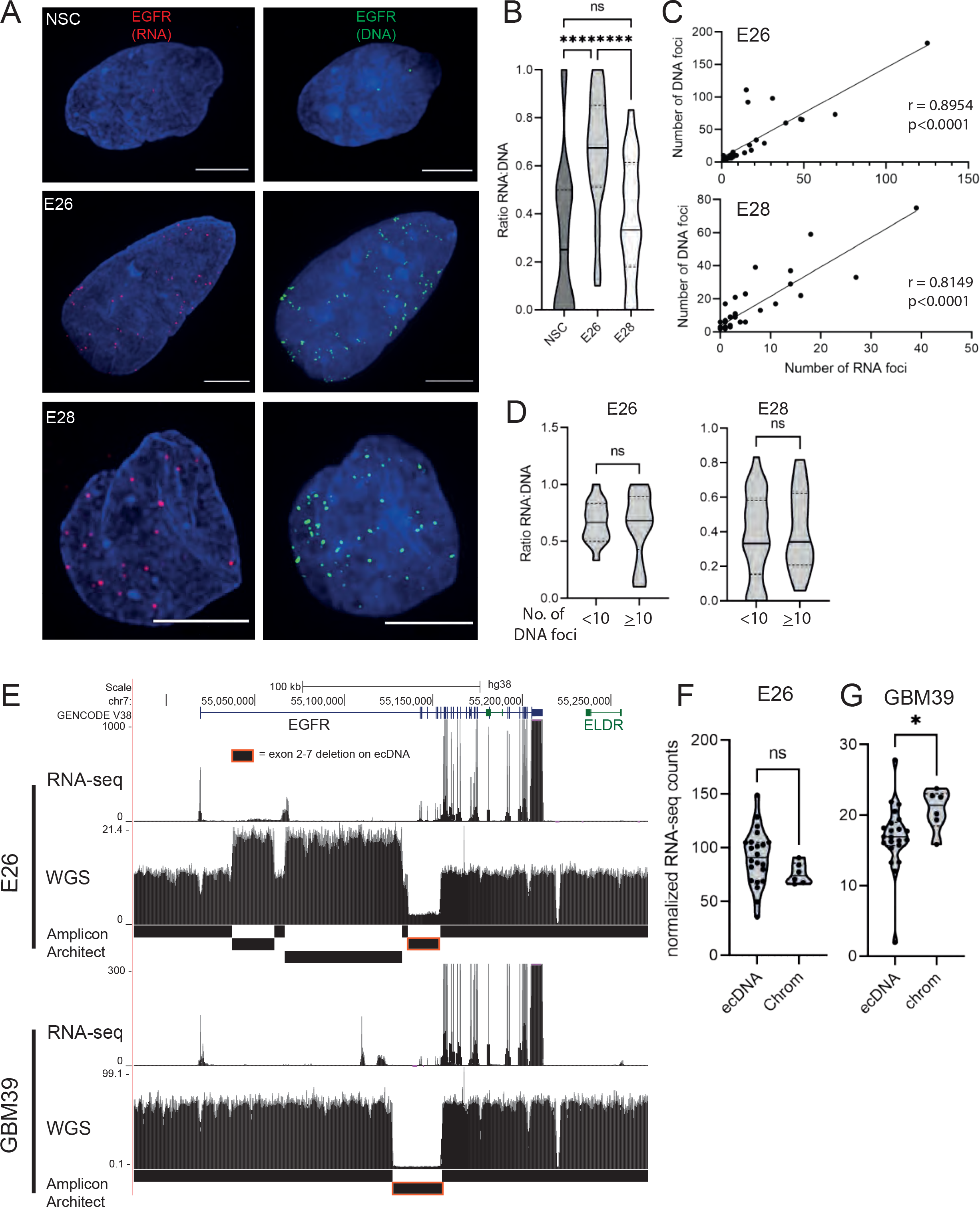
Levels of transcription from ecDNA reflect copy number but not enhanced transcriptional efficiency. A) Representative maximum intensity projection (MIP) images of nascent EGFR RNA and EGFR DNA FISH in NSC, E26 and E28 cell lines (Scale bar = 5um). B) Violin plot of ratio of RNA:DNA foci per nucleus in NSC, E26 and E28 cell lines. **** p<0.0001. C) Spearman r correlation (ρ) and p values shown for E26 and E28 cell lines. At least 33 nuclei of each cell line. D) Violin plot of ratio of RNA:DNA foci per nucleus, nuclei categorised to primarily chromosomal EGFR (<10 EGFR DNA foci) and primarily ecDNA EGFR (>10 EGFR DNA foci). ns, not significant. E) UCSC genome browser tracks showing E26 and GBM39 RNA-seq and WGS aligned sequences in the region of chromosome 7 where EGFR is located, EGFR exons (GENCODE) and the exon deletion predicted by AmpliconArchitect. Note that RNA-seq counts in some ecDNA regions go above the maximum value. Genome co-ordinates (Mb) are from the hg38 assembly of the human genome. F) EGFR RNA-seq counts in E26 and GBM39 normalized by exon size for each of the exons was normalized to the copy number (WGS counts per region normalized by region size) in regions defined by AmpliconArchitect, and labelled as extrachromosomal or chromosomal based on copy number. Statistical significance examined by Mann-Whitney test. ns, not significant, * p<0.05. Statistical data relevant for this figure are in Table S5.

To test this using an independent method, we utilized the large exon 2-7 deletion (Figure S1A) present on E26 *EGFR* ecDNA to quantify transcriptional efficiency using analysis of RNA-seq and WGS data. We compared the copy number-normalized RNA expression of exons present either only on the endogenous chromosomal *EGFR* locus (exons 2-7) with those predominantly on ecDNA (exons 1, 8-28), using AmpliconArchitect to define chromosomal and ecDNA regions (Figure 5E). Copy number and exon size-normalized EGFR RNA counts were not significantly different between exons 2-7 and those located predominantly on ecDNA (Figure 5F). EcDNA with *EGFR* in another established GBM cell line, GBM39, also contains a deletion spanning exons 2-7. We therefore repeated this analysis using previously published WGS and RNA-seq data and chromosome/ecDNA regions reported for this cell line via AmpliconArchitect (Wu et al., 2019). The normalized RNA of ecEGFR exons was not higher than that of chromosomal EGFR exons and was in fact moderately lower in GBM39 (Figure 5F). Altogether, RNA:DNA FISH imaging and sequencing analysis suggest that *EGFR* on ecDNA is transcribed at a similar level to that of the corresponding endogenous chromosomal *EGFR* locus. Increased output of oncogenes in GBM with ecDNA may primarily be driven by increased copy number, rather than inherent features of their chromatin state, transcriptional control, or spatial localization.

## Discussion

Understanding the importance of ecDNA in the etiology of cancer, and whether this poses an interesting target for therapeutic interventions, depends on deeper analysis of ecDNA activity (Kim et al., 2020; Nathanson et al., 2014). Clustering of ecDNA into ‘ecDNA hubs’ based on imaging and chromosome conformation capture data has been reported in a range of established cancer cell lines, and may therefore underlie the ability of ecDNA to drive very high levels of transcription (Hung et al., 2021; Yi et al., 2021; Zhu et al., 2021). However, in multiple primary human GBM cells studied here, we observe that while there are broader regional biases in the localization of ecDNA in the nucleus, there was no colocalization at distances close enough (∼200nm) thought to be functionally important in driving transcription.

We reach this conclusion for both cells with single ecDNA species, as well as with heterogeneous ecDNA harboring different oncogenes. EcDNA were not colocalized with, or notably close to, large PolII condensates. Moreover, taking advantage of the unique transcripts from ecDNA, we demonstrate that increased transcription of ecDNA-located genes is primarily driven by increased copy number rather than increased transcriptional efficiency of ecDNA.

Our data suggests ecDNA are widely distributed through the nucleus, but that they have a significantly different nuclear distribution than the corresponding endogenous gene loci. Chromosomes are not randomly distributed, and occupy specific nuclear territories (Boyle et al., 2001; Croft et al., 1999). Freed of the constraints imposed by being part of large chromosomes, ecDNA occupy positions away from the nuclear periphery, which is consistent with an actively transcribing state. A less peripheral nuclear localization of ecDNA is consistent with ChIA-PET data detecting transcriptionally active (RNA PolII-bound) genomic associations of ecDNA in GBM cell lines (Zhu et al., 2021). Chromosome conformation data have previously shown that ecDNA harbors highly accessible chromatin (Wu et al., 2019), which is primarily in the nuclear centre. Given that gene-rich human chromosomes (e.g., 17, 19, 22) are preferentially found toward the center of the nucleus, this region therefore represents a gene-rich, accessible environment in which ecDNA exist (Boyle et al., 2001; Croft et al., 1999). The high level of trans-interactions detected from ecDNA are also consistent with a localization outside of chromosome territories – as has been detected for highly active, decondensed, endogenous chromosomal regions that loop outside of their own chromosome territories (Kalhor et al., 2011).

In our analysis we sought to maximize our opportunity of observing ecDNA clustering at close distances by performing 3D spot analysis, using an optimistic (i.e., low) estimate of *p* values where required during Ripley’s K analysis. We also used cells with two distinct ecDNA species to ensure there was no under-scoring in the spot analysis. 3D analysis ensures a false positive clustering effect is avoided that might be seen when 3D images are combined via tools such as maximum intensity projection (MIP). Other tools to assess clustering have noted the possibility of the 2D Ripley’s K function resulting in over-counting, leading to the development of alternative auto-correlation tools (Veatch et al., 2012). We did not observe an over-counting effect given that no clustering at short distances was observed, which could be due to the 3D nature of our analysis, or the factors that lead to over-counting not being relevant to this experimental design, such as our imaging facilitating identification and resolution of individual foci. We did not observe ecDNA clustering at close distances (<300nm) in any of our cell lines in a 3D analysis, suggesting this is not a major contributor to increased ecDNA transcriptional output.

Our findings may reflect fundamentally different functional characteristics of the ecDNA in the specific patient cell cultures used in our experiments versus previously published studies (Hung et al., 2021; Yi et al., 2021). These might include the size of the ecDNA, or the number of oncogene loci per ecDNA (which was singular in our cell lines). For example, the COLO320-DM cell line, used in a recent study of ecDNA hubs, harbors 3 copies of *MYC* on each of its ecDNA, and results in large (approx. 1.75Mb) ecDNA (Hung et al., 2021; Wu et al., 2019). The HK359 GBM cell line, previously noted to have clustered ecDNA hubs, has a 42kb insertion at the site of *EGFR*vIII (exon 2-7 deletion), again suggesting a large ecDNA quite different in character to those described here (Hung et al., 2021; Koga et al., 2018). More quantitative analysis across a larger set of primary cancer cells will be needed to determine if the long term established cell lines COLO320-DM are unusual in their features and unrepresentative of primary cancer cells.

Recent work proposing that ecDNA act as mobile super-enhancers for chromosomal targets has raised the possibility that ecDNA can actively recruit RNA PolII to drive ‘ecDNA-associated phase separated condensates’ (Adelman and Martin, 2021; Zhu et al., 2021). Despite recent work in different cancer cells proposing clustering of ecDNA and RNA PolII (Yi et al., 2021), we did not observe evidence of a close relationship between ecDNA, or their nascent transcript, with PolII condensates. Although our strategy was not designed to quantify general PolII signal correlation with ecDNA – noting that granular, pan-nuclear PolII staining was seen in all imaged nuclei – our high-resolution 3D imaging approach gives us confidence that no colocalization was seen with large PolII condensates. Indeed, the low prevalence of condensates in our cell lines is in keeping with other studies (Imada et al., 2021). We also cannot exclude that there are smaller, sub diffraction-limit sized transcriptional condensates associated with our ecDNA.

We observe that while the copy number of ecDNA encoding *EGFR* positively correlate with greater transcriptional output, this is likely due to copy number increases, rather than increased transcriptional activity at individual ec*EGFR* loci. Recent work has proposed that ecDNA increase transcription of their resident oncogenes partly due to their increased DNA copy number, but also due to their more accessible chromatin structure, and that gene transcription from circular amplicons was greater than that of linear amplicons once copy number normalized (Kim et al., 2020; Wu et al., 2019). In an analysis of RNA-seq and WGS data from a cohort of 36 independent clinical samples, when normalized to gene copy number, only 3 out of 11 ecDNA-encoded genes produced significantly more transcripts, only one of which is a key oncogene, suggesting that copy number is the dominant driver (Wu et al., 2019). Transcriptional efficiency of a particular population of ecDNA in any given cell line may depend on the specific combination of regulatory elements present on the ecDNA, and there may be other genetic or epigenetic differences between established and primary cell lines that affect transcription.

Overall, our data suggest that in primary GBM cells ecDNA can succeed at driving oncogene expression without requiring close colocalization with each other, or with transcriptional condensates. It is the increased copy number that is primarily responsible for higher levels, rather than ecDNA-intrinsic features or nuclear sub-localization.

## Author contributions

K.P. conducted most of the experiments and data analysis, prepared the figures and wrote the manuscript. E.T. F. assisted in IF optimization, RNA/WGS sequencing analysis, general project design, and edited the manuscript. S.B. assisted in all aspects of DNA FISH, and general project design. P.S.D. and V.G. created and curated the E28-mCherry-POL2RG cell line. A.H. performed AmpliconArchitect analysis. G.M.M. derived and characterized the GSC primary cell lines and the NSC primary cell line. P.B. obtained the GBM primary cell tissue. S.V.B. designed and wrote the script for the Ripley’s K function analysis with KP and provided statistical oversight throughout the study. W.A.B. and S.M.P. conceived of the study, designed the study, coordinated the study, and wrote the manuscript. All the authors gave final approval for publication.

## Acknowledgements

K.P. was supported by a Wellcome PhD Training Fellowship (220399/Z/20/Z). E.T.F. was supported by the Swiss National Science Foundation (P2ELP3_191695). G.M.M. and the Glioma Cellular Genetics Resource (www.gcgr.org.uk) were supported by the Cancer Research UK (CRUK) Centre Accelerator Award (A21922). A.H. was supported by a CRUK PhD Fellowship (C157/A29279),). S.M.P is a Cancer Research UK Senior Research Fellow (A17368). Work in the group of W.A.B. is supported by MRC University Unit grant MC_UU_00007/2.

S.V.B. would like to thank Dr Tim Cannings for helpful suggestions on statistical analysis.

We acknowledge the Advanced Imaging Resource at the Institute of Genetics and Cancer and the Edinburgh Super-Resolution Imaging Consortium (ESRIC), and the Flow Cytometry team at the Centre for Regenerative Medicine, University of Edinburgh, for their technical support.

## Methods

### Lead contact

Further information and requests for resources and reagents should be directed to and will be fulfilled by the lead contacts, Wendy Bickmore and Steven Pollard.

### Materials availability

This study generated a new CRISPR engineered knock-in reporter cell line – E28 mCherry_POLR2G.

### Data and code availability

- Cell line sequencing data are available in a source manuscript pending publication and will be made publicly available as of the date of publication. Accession numbers will be listed in the key resources table
- All original code has been deposited as outlined below and is publicly available as of the date of publication.
  - Erosion Territories analysis – Code available at https://github.com/IGC-Advanced-Imaging-Resource/Purshouse2022_paper
  - Cluster analysis – Code available at https://github.com/SjoerdVBeentjes/ripleyk
- Any additional information required to reanalyze the data reported in this paper is available from the lead contact upon request.

### Experimental Model and Subject details

GSC and NSC lines from the Glioma Cellular Genetics Resource (GCGR) (http:/gcgr.org.uk) were cultured in serum-free basal DMEM/F12 medium (Sigma) supplemented with N2 and B27 (Life Technologies), 2 μg/mL laminin (Cultrex), and 10 ng/mL growth factors EGF and FGF-2 (Peprotech) (Pollard et al., 2009). Cells were split with Accutase solution (Sigma), and centrifuged approximately weekly as previously reported. All GBM cell lines were derived from treatment-naive patients, and the NSC cell line GCGR-NS9FB_B was derived from 9 week of gestation forebrain. Human GBM tissue was obtained with informed consent and ethical approval (East of Scotland Research Ethics service, REC reference 15/ES/0094). Human embryonic brain tissue was obtained with informed consent and ethical approval (South East Scotland Research Ethics Committee, REC reference 08/S1101/1).

## Method details

### Metaphase spreads and interphase nuclei

Cell lines were optimized to generate metaphase spreads. Briefly, cells at near confluence were incubated between 4 and 16 hours in the presence of 10-100nm paclitaxel (Cambridge BioScience) with or without 50-100ng/ml nocodazole (Sigma-Aldrich). Along with the media, cells dissociated with accutase were centrifuged, washed in PBS, and resuspended in 10ml potassium chloride (KCl) 0.56%, with sodium citrate dihydrate (0.9%) if required, for 20 min. After further centrifugation, cells were resuspended in methanol:acetic acid 3:1 and dropped onto humidified slides.

For all other fixed cell experiments described below, cells were seeded overnight onto glass cover-slips or poly-L-lysine coated glass slides (Sigma-Aldrich). Cells were fixed with 4% paraformaldehyde (PFA – 10 mins) and permeabilized with 0.5%Triton X100 (15 mins) with thorough PBS washes in between.

### DNA FISH

A detailed method for DNA FISH has been described elsewhere (Jubb and Boyle, 2020). Briefly, DNA stocks of fosmid clones targeting EGFR (WI2-2910M03), CDK4 (WI2-0793J08) and PDGFRA (WI2-2022O22) (Table S6) were prepared via an alkaline lysis miniprep protocol (Jubb and Boyle, 2020). Each fosmid DNA probe was labelled via Nick Translation directly to a fluorescent dUTP (Green496-dUTP, ENZO life sciences; ChromaTide AlexaFluor 594-5-dUTP, ThermoFisher Scientific) and incubated with unlabelled dATP, dCTP and dGTP, ice cold DNase and DNA polymerase I for 90 min at 16°C. The reaction was quenched with EDTA and 20% SDS, TE buffer added and the reaction mix run through a Quick Spin Sephadex G50 column.

Cells on slides or cover-slips were prepared by incubating for one hour in 2x Trisodium citrate and sodium chloride (SSC)/RNaseA 100 μg/ml at 37°C, then dehydrated in 70%, 90% and 100% ethanol. Slides were warmed at 70°C prior to immersion in a denaturing solution (2xSSC/70% formamide, pH 7.5) heated to 70°C (methanol:acetic acid-fixed cells) or 80°C (PFA-fixed cells), the duration of which was optimized to each cell line. After denaturing, slides were immersed in ice-cold 70% ethanol, then 90% and 100% ethanol at room temperature before air drying.

FISH probes were prepared by combining 100ng of each directly labelled fosmid probe (per slide), 6 μg Human Cot-1 DNA (per probe), 5 μg sonicated salmon sperm (per slide) and 100% ethanol. Once completely dried, the resulting pellet was suspended in hybridization mix (50% deionized formamide (DF), 2× SSC, 10% dextran sulfate, 1% Tween 20) for one hour at room temperature, denatured for 5 mins at >70°C and annealed at 37°C for 15 min. Where relevant, FISH probes were instead hybridized in Chromosome 7 paint (XCP 7 Orange, Metasystems). The probes were incubated overnight at 37°C. The following day, the slides were washed in 2x SSC (45°C), 0.1% SSC (60°C) and finally in 4xSSC/0.1% Tween 20 with 50ng/ml 4′,6-diamidino-2-phenylindole (DAPI). Slides were mounted with Vectashield.

### RNA FISH

RNA FISH probes (Custom Assay with Quasar® 570 Dye) targeting the first intron (pool of 48 22-mer probes) of *EGFR* were designed and ordered via the Stellaris probe designer (Biosearch Technologies, Inc., Petaluma, CA) (https://www.biosearchtech.com/support/tools/design-software/stellaris-probe-designer, version 4.2). Cells were seeded, fixed and permeabilized as above. Slides were immersed in 2xSSC, 10% DF in DEPC treated water for 2-5 min before applying the hybridization mix (Stellaris RNA FISH hyb buffer, 10% DF, 125nm RNA FISH probe) for overnight incubation at 37°C. 24hr later, slides were incubated in 2xSSC, 10% formamide in DEPC-treated water for 30 min, and then repeated with DAPI (5ng/ml). Slides were washed with PBS before mounting with Vectashield.

For combined RNA-DNA FISH, nascent EGFR RNA FISH was performed as above, and nuclei imaged as described below. The x,y,z coordinates for each image were recorded via NIS software at the time of imaging. After removing the cover slips and washing the slides in PBS, EGFR DNA FISH was performed whereby the probe preparation was as above. Slides were transferred from PBS wash to denaturing solution at 80°C for 15 min, washed in 2xSSC, and incubated overnight with the probe at 37°C. The subsequent wash and mounting steps were as described above. The stored x,y,z coordinates were used to relocate and image each nucleus. Owing to the irregularity of the tumor nuclei, it was possible to be confident in re-imaging the correct nucleus. Spot counting was subsequently performed as described below with RNA and DNA foci being defined and counted separately to avoid influencing the outcome.

### Immunofluorescence and immuno-FISH

Slides were blocked in 1%BSA/PBS/Triton X-100 0.1% for 30 min at 37°C before overnight incubation with the primary antibody at 4°C (Rpb1 NTD (D8L4Y) #14958, Cell Signalling Technology, 1 in 1000; mCherry (ab167453), abcam, 1 in 500). The following day, slides were washed in PBS before incubation with an appropriate secondary antibody (1:1000 AlexaFluor) for one hour at 37°C. After further PBS washes and DAPI staining, slides were mounted with Vectashield.

For immuno-FISH (DNA), the IF signal was fixed via incubation with 4% PFA for 30 minutes. Following thorough PBS washes, the DNA FISH protocol was then followed as above.

For immuno-FISH (RNA), the antibodies were added at the same concentration as described above to the hybridization mix (primary antibody) and 2xSSC/10% DF washes (secondary antibody).

### Flow Cytometry and FACS

Cells were prepared by adding EGF-free media for 30 minutes before lifting and suspending cells in 0.1% BSA/PBS. Cells were incubated in 100ng/ml EGF-647 (E35351, Thermo Fisher Scientific) in 0.1%BSA/PBS, with cells incubated in 0.1% BSA/PBS as a negative control, for 25 minutes. Cells were washed three times in 0.1%BSA/PBS before being analyzed on the BD FACSAria III FUSION. Where indicated, cells were sorted by EGF-647 gated into high and low groups, and a sort check was performed to verify these were true populations prior to expanding these cells onto 22×22mm cover slips. Fifteen days after the cells were sorted, the slides were fixed, permeabilized and DNA FISH performed as above.

### mCherry_PolR2G knock in cell line

crRNA and donor DNA was designed using the previously reported TAG-IN tool (Dewari et al., 2018), with the corresponding fluorescent reporter gene sequences for mCherry implemented into the existing tool (Table S7). Output sequences from the TAG-IN tool were manufactured by Twist Bioscience. Gene-specific crRNA (100 pmoles - IDT Technologies, USA) and universal tracrRNA (100 pmoles, IDT Technologies, USA) were assembled to a cr:tracrRNA complex by annealing at the following settings on a PCR block: 95°C for 5 min, step down cooling from 95°C to 85 °C at 0.5°C/sec, step down cooling from 85°C to 20°C at 0.1°C /sec, store at 4°C. Recombinant Cas9 protein (10μg, purified in house - see (Dewari et al., 2018)) were added to form the ribonucleoprotein (RNP) complex at room temperature for 10 minutes, then stored on ice. 300ng of donor dsDNA were denatured in 30% DMSO by incubating at 95 °C for 5 min followed by immediate immersion in ice. The donor dsDNA and RNPs were electroporated into E28 cells using the 4D Amaxa X Unit (programme DN-100). After two weeks of serial expansion of cells in 2D culture, assessment of knock-in efficiency was assessed by suspending 5-7x 10^5^ cells in 0.2% BSA/PBS and analysed on BD LSRFortessa™ Cell Analyzer, with cells electroporated with tracrRNA:Cas9 only as a negative control. Cells were then further sorted into a pure KI population, and mCherry KI was verified by immunofluorescence for mCherry and Rpb1.

### Imaging

Slides were imaged on epifluorescence microscopes (Zeiss AxioImager 2 and Zeiss AxioImager.A1) and the SoRa spinning disk confocal microscope (Nikon CSU-W1 SoRa). For 3D image analysis, images were taken with the SoRa microscope and a 3 μm section across each nucleus was imaged in 0.1 μm steps. Images were denoised and deconvolved using NIS deconvolution software (blind preset) (Nikon). 3D images are shown in the figures as maximum intensity projections (MIP) prepared using ImageJ.

### Quantification and Statistical Analysis

#### Image analysis of nuclear localization

Images were analysed using Imarisv9.7 and Fiji. The scripts used to perform nuclear territory analysis have been described elsewhere (Boyle et al., 2001; Croft et al., 1999). Briefly, single-slice images were taken with a 20x lens using the Zeiss AxioImager 2, imaging at least 50 nuclei per cell line. The images were segmented first to individual nuclei, and subsequently the area of the DAPI signal was segmented to define the nuclear area. This area was segmented into concentric shells of equal area from the periphery to the centre of each nucleus. The signal intensity of each FISH probe or chromosome paint signal was calculated, with normalisation for the DAPI signal in each shell.

#### Image analysis of ecDNA and condensates

For 3D analysis, deconvolved images were analysed using Imaris (v9.7) and all analysis was performed on the full 3D image. RNA and DNA FISH foci, and where relevant, condensates, were defined, counted and distances between them calculated, using the Spots function within Imaris. In all cases, the Spots function was used to find the shortest distance between the objects in question.

For 3D cluster analysis of FISH spots, Ripley’s K function was performed using the x,y,z coordinates for each FISH spot using the Imaris Spots function to determine observed and null distribution values.

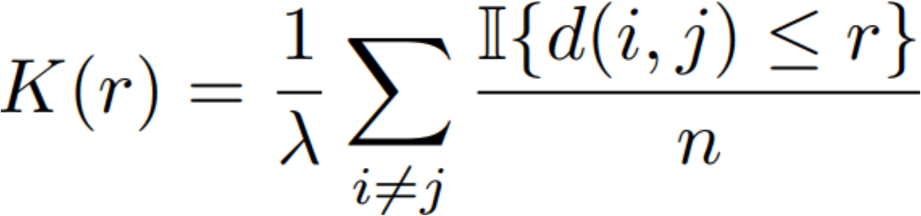

Ripley’s K function compares the number of points at a distance smaller than a given radius r, relative to the average number of points in the volume. This average is the density lambda, in this case the number of foci, n, divided by the volume. In the above equation, II is the indicator function which equals one if the distance between points i and j is no larger than r, and zero otherwise. A high value of Ripley’s K function represents clustering at the given radius r, whereas a low value represents dispersion. Consequently, a high Ripley’s K function at a given radius is indicative of clustering at this radius. By comparing the observed value of Ripley’s K function at a given radius with that computed on the same number of foci and with the same volume but drawn from a uniform null distribution, the presence of significant clustering in the given cluster at the given radius can be detected.

The code written to perform this analysis was formed using a script written in Python (v3.9) and has been made available on Github (See Data and Code Availability above). Ripley’s K function was determined across a radius of 0.1-1um in 0.1um increments. After calculating the observed Ripley’s K function value, a null distribution of no clustering, estimated on uniformly distributed samples with the same number of spots, was generated using the coordinates for each given nucleus to calculate 10,000 Ripley’s K function values at each radial increment. We tested a sample of nuclei with 50,000 values and confirmed that 10,000 values would provide sufficient accuracy. Having sampled that nucleus shape and size did not affect the significance of a result at each increment in the given range of radii, a bounding radius of 5 was used for all samples. Only nuclei with greater than 20 *EGFR* foci were included to ensure both that the majority of foci were ec*EGFR*, to allow adequate granularity and minimize the risk of a false negative result due to lack of foci. The p-value for each observed K-function was established against the expected values using the Neyman-Pearson lemma. Where the observed and expected K-function at p=0.05 were the same, a randomized binomial test was performed to determine if p<0.05 for the observed value, weighting the probability of success as the ratio of the number of values p<0.05 and the total number of equal values. Having determined this, the most optimistic estimate of p-value was made which would favor identification of a significant result i.e., a bias in favor of significant clustering. A Benjamini-Hochberg Procedure was performed to control for the false discovery rate (FDR = 0.05).

All other statistical analysis was performed with Graphpad Prism v9.0. The statistical details for each experiment can be found in the relevant figure legends and in the Supplementary Tables. For figures, p values are represented as follows: * < 0.05, ** < 0.01, *** < 0.001, **** < 0.0001. Where appropriate, Bonferroni correction for multiple hypothesis testing was performed, and, where relevant, corrected p values are those plotted in the figures and are given in the Supplementary Tables.

#### RNA and WGS sequencing sample preparation, analysis and processing

The preparation of these cell lines for RNA-seq has been described in detail elsewhere (Gangoso et al., 2021). WGS was undertaken by BGI Tech Solutions with PE100 and normal library construction. WGS, RNA-seq and AmpliconArchitect data for GBM39 was taken from data made available via publication and in the NCBI Sequence Read Archive (BioProject: PRJNA506071) (Wu et al., 2019). Sequences were aligned to hg38 with STAR 2.7.1a with settings ‘--outFilterMultimapNmax 1’ used for WGS and RNA-seq data and settings ‘--alignMatesGapMax 2000 --alignIntronMax 1 --alignEndsType EndToEnd’ used only for WGS data (Dobin et al., 2013). Duplicate reads were removed using Picard (Broad Institute). AmpliconArchitect (Deshpande et al., 2019) and AmpliconClassifier (Kim et al., 2020) were used to predict the ecDNA regions and classify circular amplicons for E26. Exon coordinates were extracted from Ensembl (isoform:EGFR-201, Ensembl Transcript ID: ENST00000275493.7). Alignments were converted to bigWig files using deepTools bamCoverage with setting ‘--normalizeUsingRPKM’ (Ramírez et al., 2016). HOMER2 (Heinz et al., 2010) makeTagDirectory and annotatePeaks.pl (settings ‘-len 0 -size given’) were used for read counting of WGS within the ecDNA/chromosome blocks defined by AmpliconArchitect, and RNA in EGFR exons. Analysis of RNA-seq counts per copy number was performed using a script written in Python (v3.9). WGS and RNA-seq read counts were normalised to the size of the ecDNA/chromosome blocks and EGFR exons, respectively. Normalized RNA-seq read counts of each exon were divided by the normalised WGS read counts of the corresponding ecDNA/chromosome region to give a normalized RNA-seq count for each exon, and analysed in Graphpad Prism v9.0.

### Key Resources

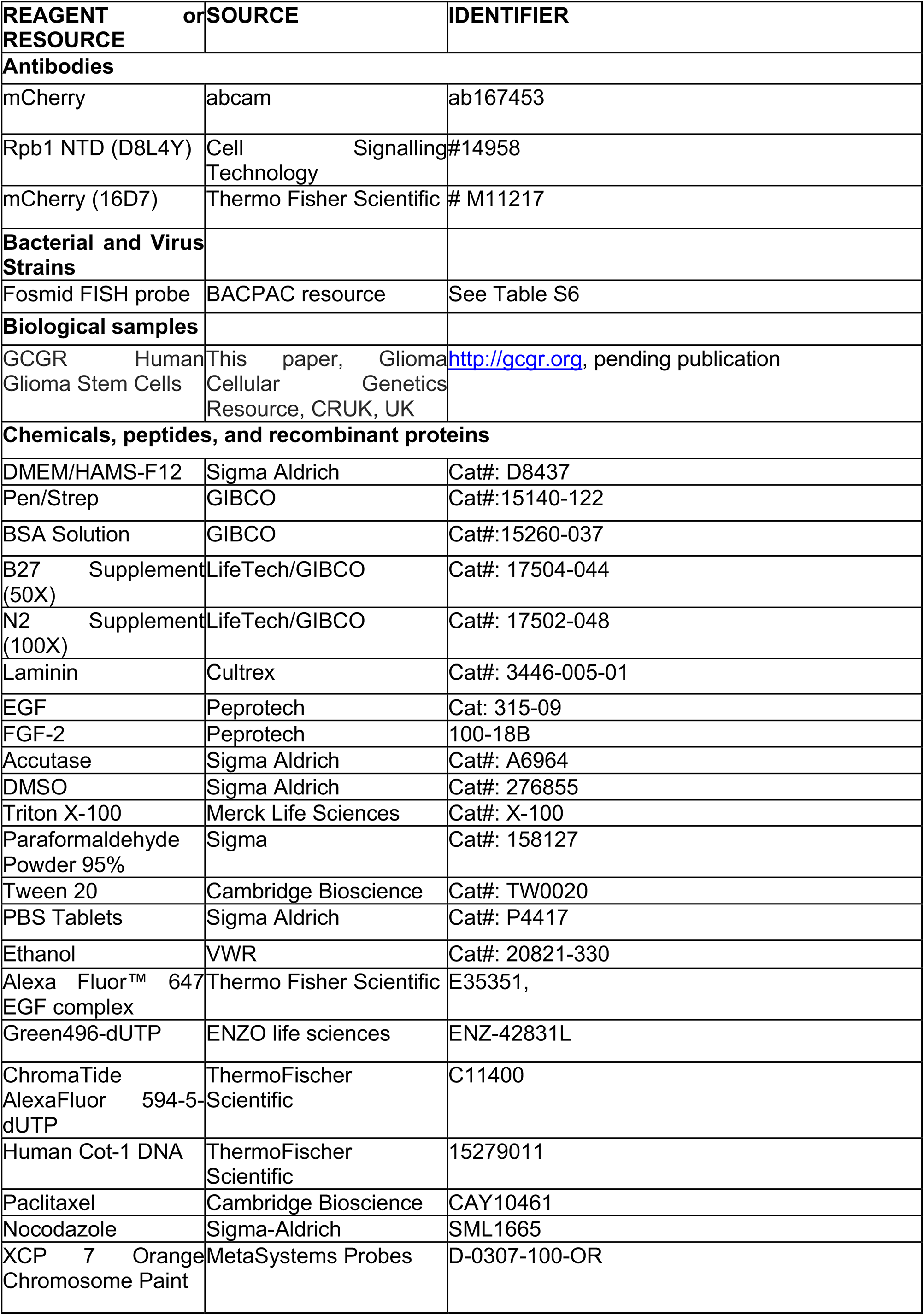
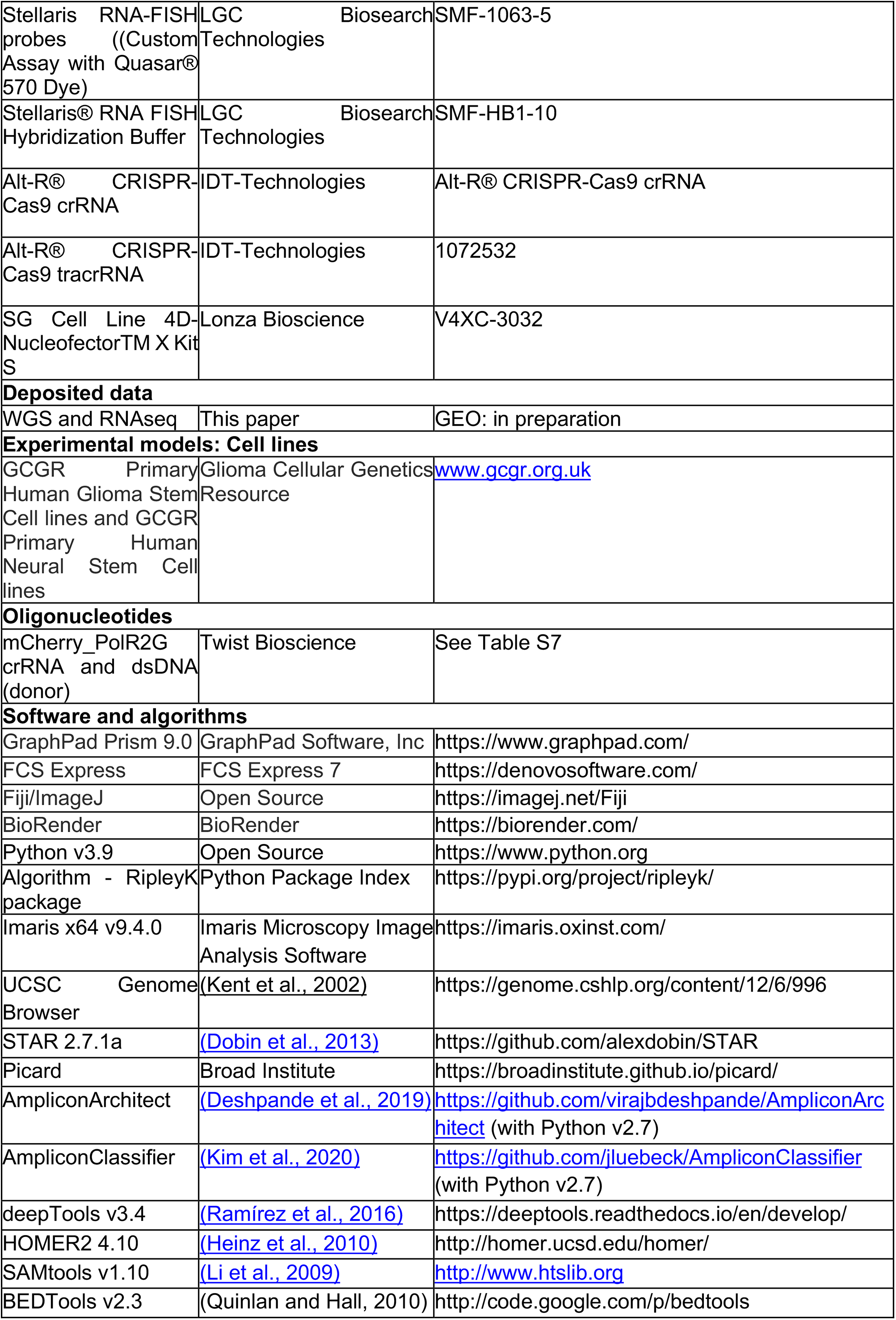

## Supplementary Information

### Supplementary Tables

**Table S1.** The nuclear localisation of ecDNA in GBM cell lines.

A. Number of EGFR DNA FISH foci per metaphase spread
|  | Cell line |  | Mann-Whitney test |
| --- | --- | --- | --- |
| Fosmid | **E26** | **E28** |  |
| **EGFR** | 51 | 12 | p = 0.0078 |

B. EGFR FISH signal intensity
|  |  | Bin |  |  |  |  | Kruskal-Wallis |
| --- | --- | --- | --- | --- | --- | --- | --- |
| Cell line | Signal | 1 | 2 | 3 | 4 | 5 |  |
| **NSC** | FITC:TxR | 0.5541 | 0.4389 | 0.3918 | 0.3756 | 0.3867 | p <0.0001 |
| **E26** | FITC:TxR | 0.6095 | 0.5573 | 0.5715 | 0.6239 | 0.6618 | p = 0.0598 |
| **E28** | FITC:TxR | 0.4802 | 0.4208 | 0.4105 | 0.4207 | 0.4682 | p = 0.0117 |
| **NSC** | TxR | 0.6159 | 0.5656 | 0.5771 | 0.5399 | 0.4732 | p <0.0001 |
| **E26** | TxR | 0.4292 | 0.3699 | 0.3279 | 0.2979 | 0.2665 | p <0.0001 |
| **E28** | TxR | 0.6563 | 0.6023 | 0.5655 | 0.4960 | 0.4175 | p <0.0001 |
| **NSC** | FITC | 0.3301 | 0.2475 | 0.2131 | 0.1991 | 0.1972 | p <0.0001 |
| **E26** | FITC | 0.2625 | 0.2034 | 0.1838 | 0.1802 | 0.1661 | p <0.0001 |
| **E28** | FITC | 0.3235 | 0.2554 | 0.2236 | 0.2117 | 0.2048 | p <0.0001 |

**Table S2.** EGFR-containing ecDNA do not cluster in the nucleus.

|  | Cell line |  | Mann-Whitney test |
| --- | --- | --- | --- |
|  | E26 | E28 |  |
| Mean shortest interprobe distance | 0.1513 | 2.092 | p = 0.1326 |
| Shortest interprobe distance | 0.4695 | 1.315 | p = 0.0322 |

**Table S3:**
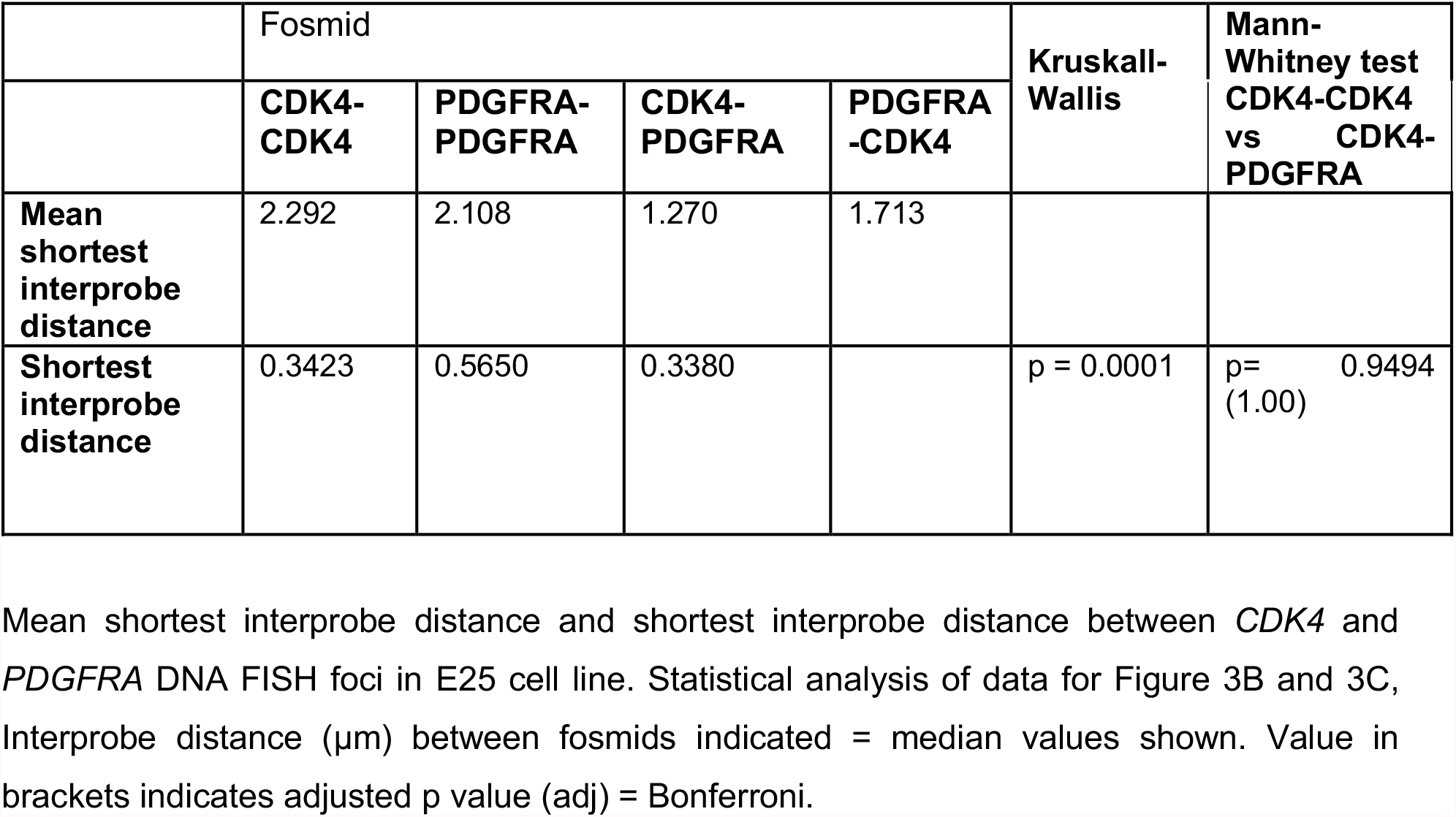
Two separate ecDNA populations do not cluster in the nucleus.

|  | Fosmid |  |  |  | Kruskal-Wallis | Mann-Whitney test<br>CDK4-CDK4<br>vs CDK4-PDGFR |
| --- | --- | --- | --- | --- | --- | --- |
|  | CDK4-CDK4 | PDGFR-PDGFR | CDK4-PDGFR | PDGFR-CDK4 |  |  |
| Mean shortest interprobe distance | 2.292 | 2.108 | 1.270 | 1.713 |  |  |
| Shortest interprobe distance | 0.3423 | 0.5650 | 0.3380 |  | p = 0.0001 | p= 0.9494<br>(1.00) |

**Table S4:** Transcriptional condensates are few and do not colocalize with ecDNA.

**A. Number of RPB1 condensates**
|  | Cell line |  | Unpaired T test |
| --- | --- | --- | --- |
|  | E26 | E28 |  |
| **Number of Rpb1 condensates** | 0.4923 (0.9206) | 0.8615 (1.321) | p < 0.0001 |

**B. ecDNA-condensate distances**
|  | Cell line |  |  | Kruskall-Wallis | Mann-Whitney test |
| --- | --- | --- | --- | --- | --- |
|  | NSC | E26 | E28 |  |  |
| **Mean shortest ecDNA-condensate distance (EGFR DNA-Rpb1)** | 3.950 | 2.995 | 1.970 | p=0.1269 |  |
| **Shortest ecDNA-condensate distance (EGFR DNA-Rpb1)** | 2.920 | 2.600 | 1.900 | p=0.5234 |  |
| **Mean shortest ecDNA-condensate distance (EGFR RNA-Rpb1)** |  | 4.667 | 5.008 |  | p=0.6277 |
| **Shortest ecDNA-condensate distance (EGFR RNA-Rpb1)** |  | 1.766 | 2.437 |  | p=0.1802 |

**C. ecDNA-condensate distances**
|  | Cell line |
| --- | --- |
|  | **E28 mCherry-PolR2G** |
| **Mean shortest ecDNA-condensate distance** | 4.804 |
| **Shortest ecDNA-condensate distance** | 1.561 |

**Table S5:** Levels of transcription from ecDNA reflect copy number but not enhanced transcriptional efficiency.

A. RNA:DNA FISH EGFR foci ratios
|  | RNA:DNA ratio | Mann-Whitney test vs E26 | Mann-Whitney test vs E28 |
| --- | --- | --- | --- |
| **NSC** | 0.2500 | p < 0.0001 | p = 0.1978 (0.5934) |
| **E26** | 0.6749 |  |  |
| **E28** | 0.3333 | p < 0.0001 |  |

B. RNA:DNA FISH EGFR foci ratios according to predominantly chromosomal (<10) vs extra-chromosomal (≥10) EGFR DNA foci.
|  | RNA:DNA ratio |  | Mann-Whitney test |
| --- | --- | --- | --- |
|  | <10 EGFR DNA foci | ≥10 EGFR DNA foci |  |
| **E26** | 0.6667 | 0.6831 | p = 0.7870 |
| **E28** | 0.3333 | 0.3417 | p = 0.8559 |

C. Normalised RNA-seq counts
|  | Normalised RNA-seq counts |  | Mann-Whitney test |
| --- | --- | --- | --- |
|  | ecDNA (n=22) | Chromosomal (n=6) |  |
| **E26** | 90.89 | 74.12 | p = 0.1122 |
| **GBM39** | 16.93 | 21.38 | p = 0.0331 |

D. EGFR foci counts
|  | Number of EGFR foci |  | Mann-Whitney test |
| --- | --- | --- | --- |
|  | EGFR high | EGFR low |  |
| **E26** | 58 | 8 | p < 0.0001 |
| **E26** | 11 | 3 | p < 0.0001 |

E. Nascent EGFR RNA foci counts
|  | Number of EGFR RNA foci | Mann-Whitney test (vs E26) | Mann-Whitney test (vs E28) |
| --- | --- | --- | --- |
| **NSC** | 1.0 | p < 0.0001 | p = 0.0174 (0.0522) |
| **E26** | 5.0 |  |  |
| **E28** | 2.0 | p = 0.0010 (0.003) |  |

|  | RNA:DNA ratio |  | Mann-Whitney test |
| --- | --- | --- | --- |
| | <6 EGFR DNA foci | $\geq 6$ EGFR DNA foci | |
| **E26** | 0.5000 | 0.3192 | p = 0.7409 |

**Table S6:** Fosmid probes for DNA FISH.

| Locus | Gene | Fosmid probes |  |  |  |
| --- | --- | --- | --- | --- | --- |
|  |  | Start (bp) | End (bp) | Fosmid ID | Clone name |
| 7: 55019017-55211628 | EGFR | 55024189 | 55063180 | G248P88704G2 | WI2-2910M03 |
| 12: 57747727-57752310 | CDK4 | 57746054 | 57783419 | G248P80931E4 | WI2-0793J08 |
| 4: 54,229,293-54298245 | PDGFRA | 54230802 | 54268615 | G248P86466H11 | WI2-2022O22 |

Related to STAR Methods. Genome co-ordinates (Mb) are from the hg38 assembly of the human genome.

**Table S7:**
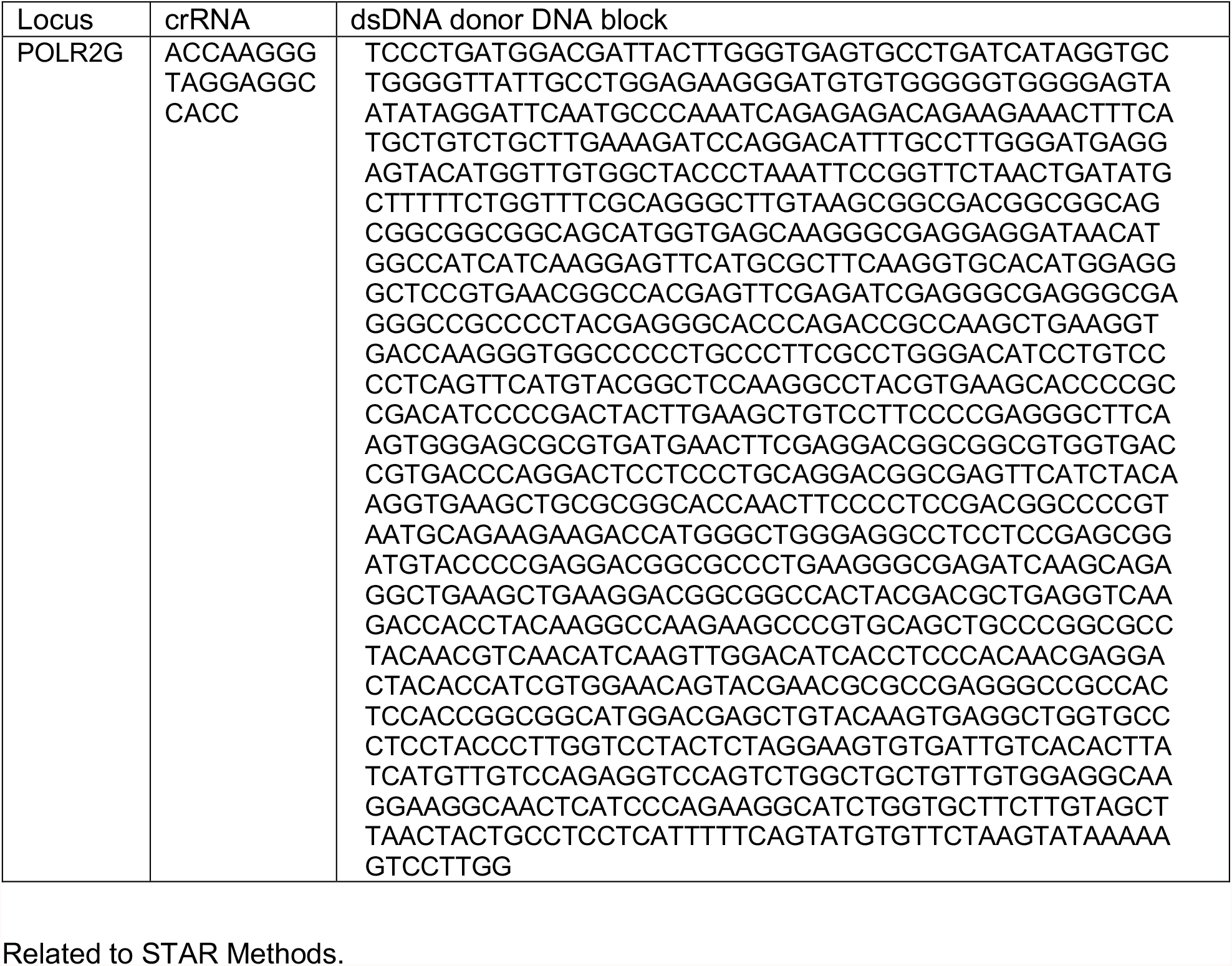
CrRNA sequence and dsDNA sequence for mCherry_PolR2G CRISPR knock-in.

**Figure S1.**
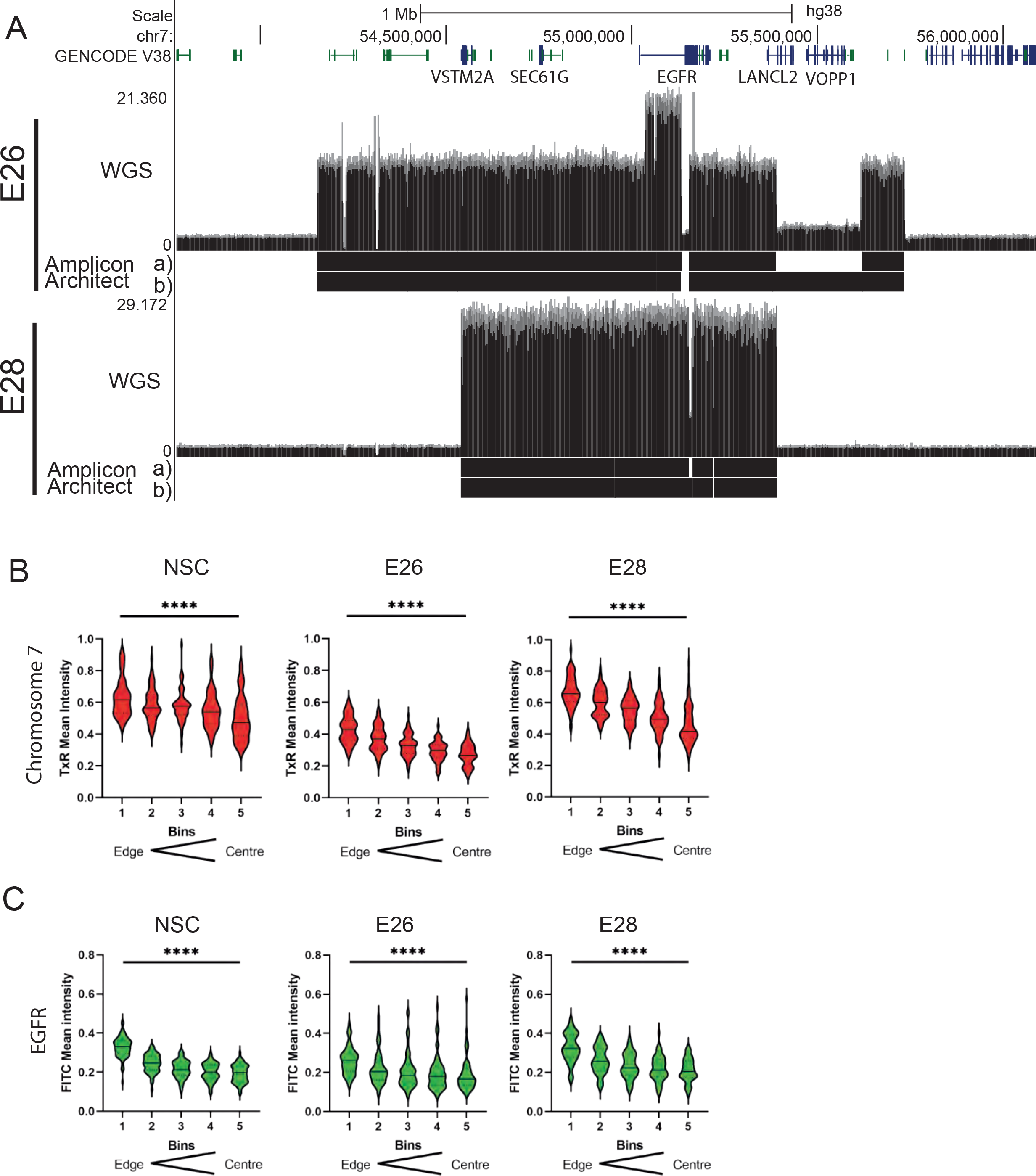
Properties of ecDNA in GBM cell lines. A) WGS and two AmpliconArchitect ecDNA regions for E26 and E28 cell lines showing an EGFR exon 2-7 deletion in all ecDNA in E26 cells (seen in WGS and AmpliconArchitect regions a and b), and a subpopulation of ecDNA in E28 with a deletion across EGFR exons 7-14 (seen in WGS and Amplicon Architect region a – no deletion in E28 AmpliconArchitect region b). Genome co-ordinates (Mb) are from the hg38 assembly of the human genome. B and C) Radial distribution, normalised to DAPI, across bins of equal area eroded from the edge (1) to the centre (5) of the nucleus for (B) Chromosome 7 (TxR Mean normalised Intensity per nucleus) or (C) EGFR (FITC Mean normalised Intensity per nucleus). Median and quartiles are shown. Statistical significance was examined by Kruskall-Wallis. **** p<0.0001. Statistical data relevant for this figure are in Table S1B.

**Figure S2.**
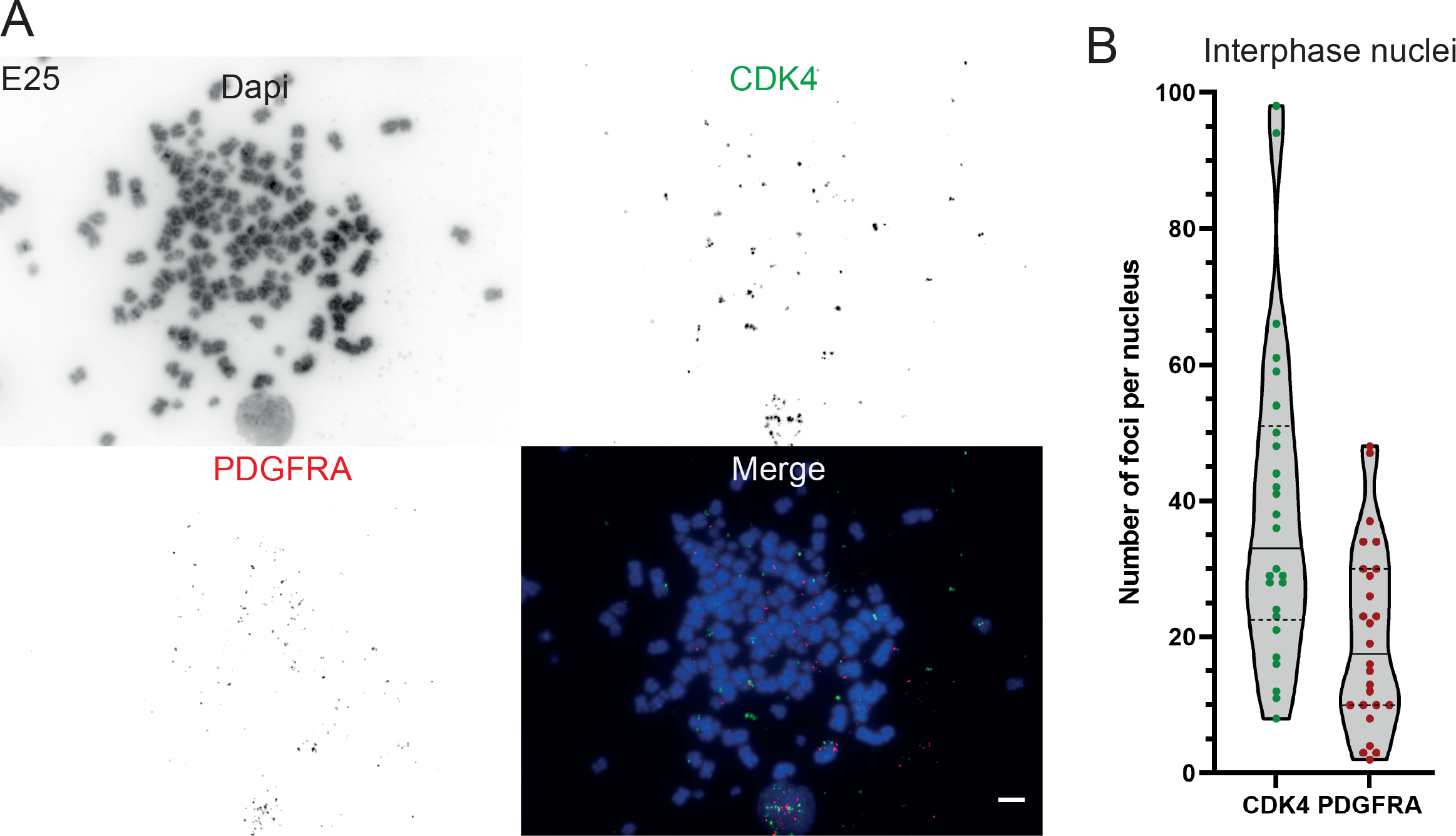
Characterisation of two ecDNA populations in E25 cells. A) DNA FISH for CDK4 and PDGFRA on metaphase spreads from the E25 cell line, showing CDK4 and PDGFRA on separate ecDNA, scale bar = 5 μm. B) Number of ecDNA per nucleus in E25 cell line, CDK4 (green) and PDGFRA (red). Number of nuclei = 26.

**Figure S3.**
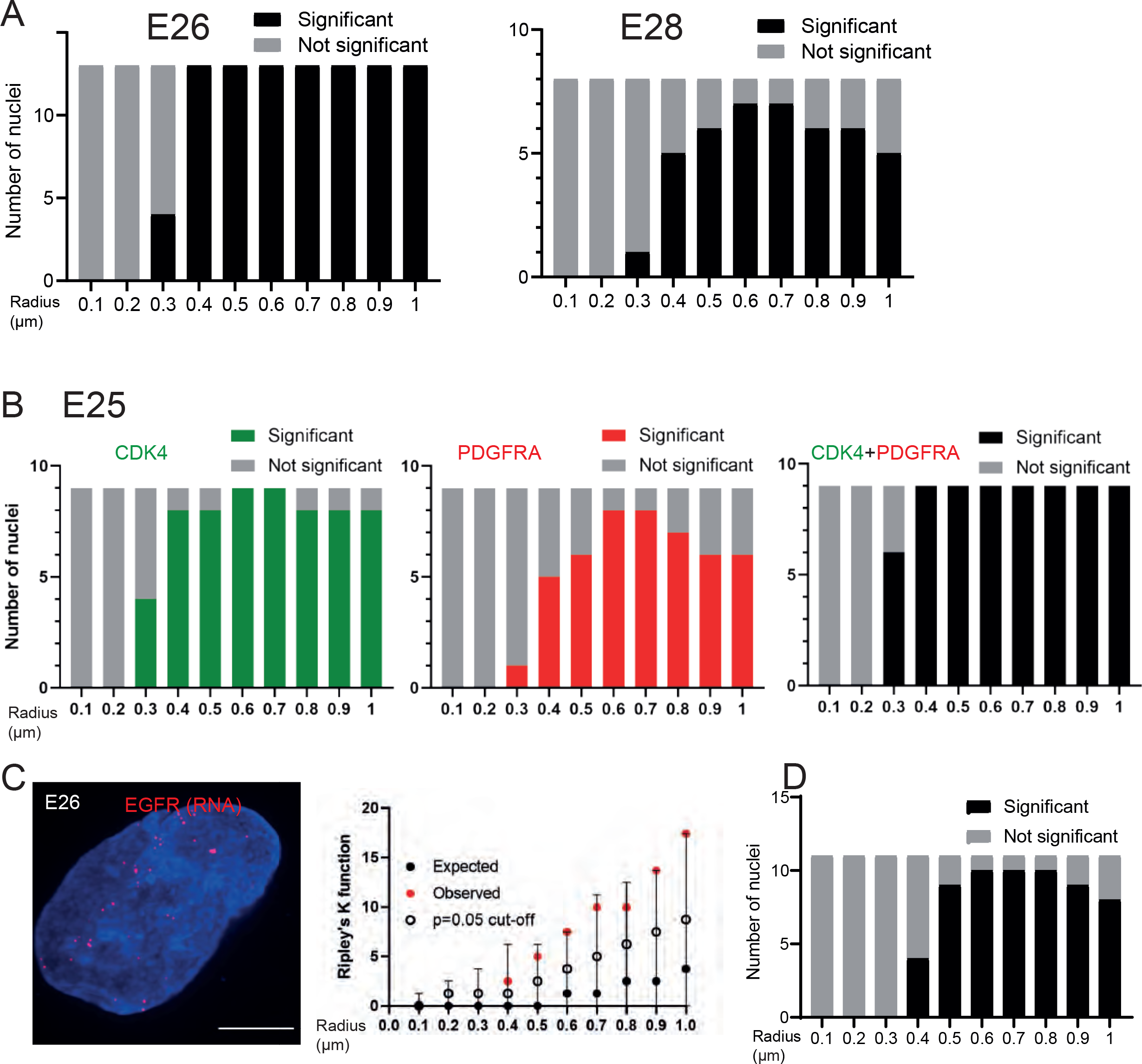
Ripley’s K function of ecDNA distribution in E26, E28 and E25 cell lines. A) Ripley’s K function for E26 and E28 nuclei showing number of nuclei with significant and non-significant clustering at each given radius - with spot diameter = 150nm. B) Ripley’s K function for E25 nuclei showing number of nuclei with significant and non-significant clustering at each given radius for CDK4 spots, PDGFRA spots and CDK4 and PDGFRA spots combined - with spot diameter = 150nm. C) Representative image shown as MIP of E26 nascent EGFR RNA FISH, scale bar = 5 μm. Associated Ripley’s K function for this nucleus showing observed K function (red), Max/Min/Median (black) of 10,000 null samples with p=0.05 significance cut-off shown (empty black circle). D) Ripley’s K function for E26 nuclei after EGFR nascent RNA FISH showing number of nuclei with significant and non-significant clustering at each given radius. All p values calculated using Neyman-Pearson lemma with optimistic estimate p value where required, and Benjamini-Hochberg Procedure (BHP, FDR = 0.05).

**Figure S4.**
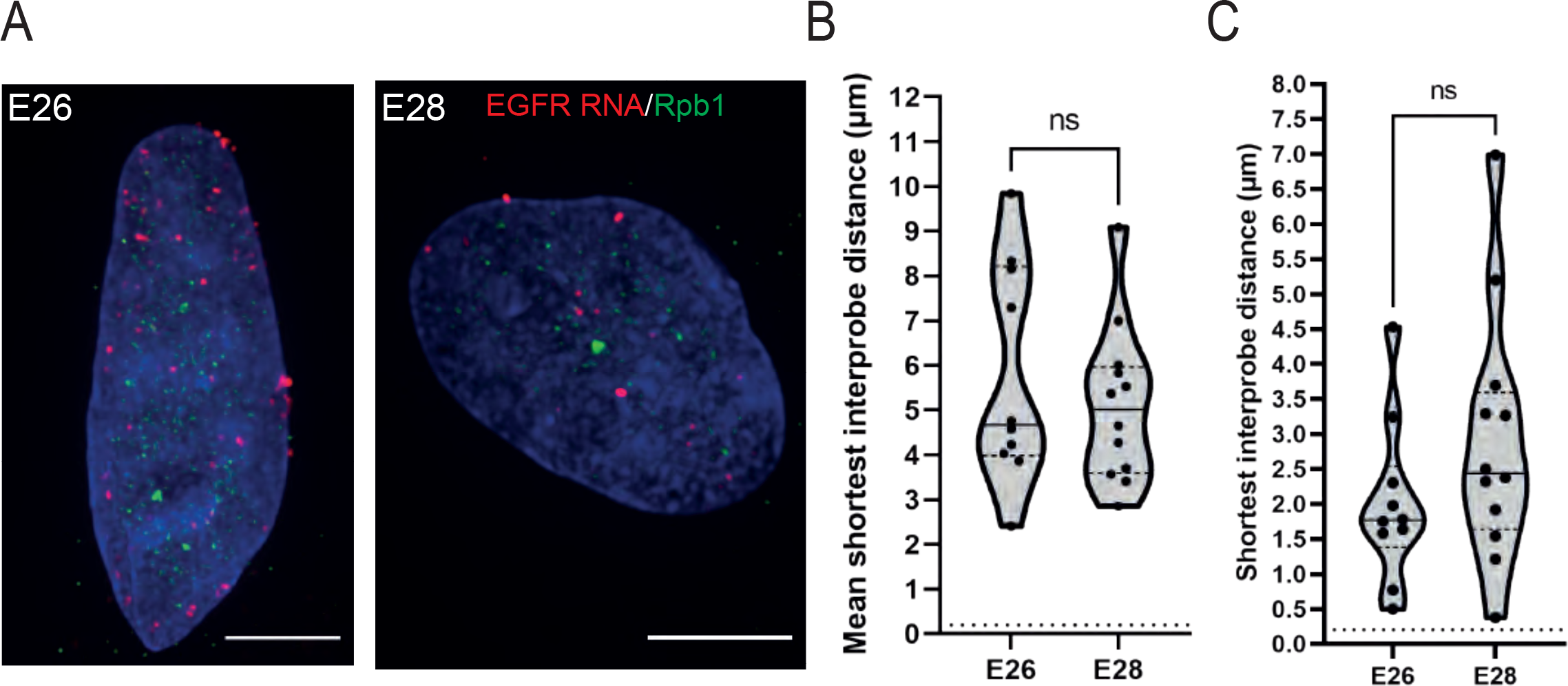
Transcribing ecDNAs do not co-localise with RNA PolII condensates. A) Representative images of E26 and E28 nascent RNA immunoFISH for EGFR (red) and RPolII (Rpb1 – green), MIP, Scale bar = 5 μm. B) Mean shortest interprobe distance between EGFR RNA foci and condensates. C) As for (B) but for shortest distance. Median and quartiles plotted. Dotted line denotes y = 200nm. Statistical significance examined by Mann-Whitney. ns, not significant. Statistical data relevant for this figure are in Table S4B.

**Figure S5:**
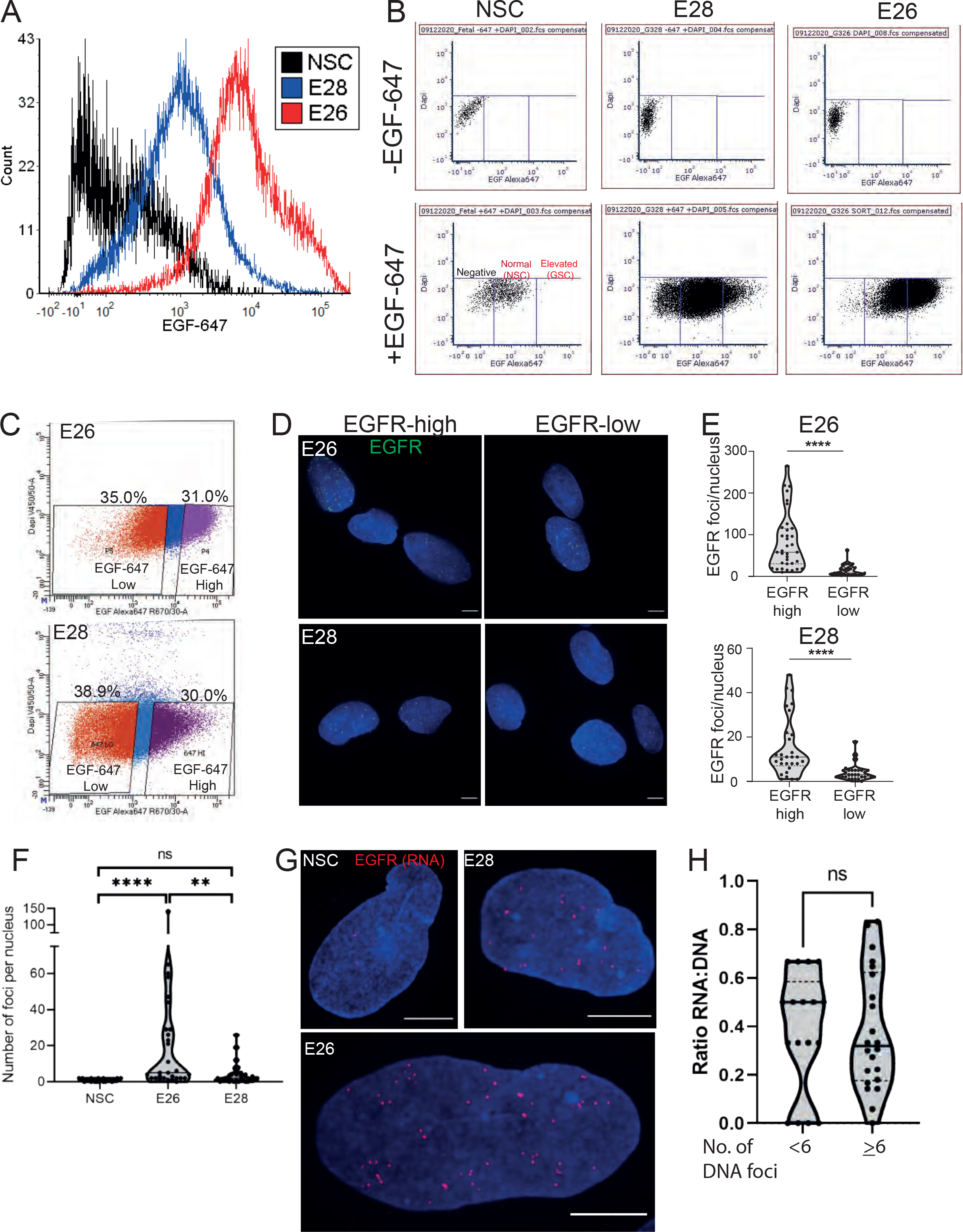
Characterisation of EGFR-high and EGFR-low cell populations. A) Histogram of flow cytometry with EGF-647 showing signal in NSC, E28 and E26 cell lines from live cells, normalised to peak count per cell line. Median EGF-647 - NSC = 172.2; E28 = 985.64; E26 = 7191.81. B) Flow cytometry with EGF-647; gates showing negative, normal (NSC) and elevated (GSC) EGF-647 signal in NSC, E28 and E26 cell lines. C) Fluorescence activated cell sorting (FACS) into EGF-647 high and low populations from E26 and E26 cell lines. The percentage of total live cell population in each sorted population are shown. D) Representative EGFR DNA FISH images of E26 and E28 cells sorted via flow cytometry with EGF-647 into EGFR high and low cells. MIP, scale bar = 5 μm. E) Number of EGFR DNA FISH per nucleus in sorted E26 and E28 cells. Statistical significance examined by Mann-Whitney. **** p<0.0001. Statistical data relevant for this figure are in Table S5D.F) Number of nascent EGFR RNA foci per cell line, at least 25 nuclei of each cell line imaged. Statistical significance examined by Mann-Whitney test. ns = not significant, ** p<0.01, ****p<0.0001 and are detailed in Table S5E. G) Representative images of nascent EGFR RNA FISH in NSC, E26 and E28 cell lines. MIP, scale bar = 5 μm. H) Ratio of RNA:DNA foci per nucleus in E28 cell line, nuclei categorised to primarily chromosomal EGFR (<6 EGFR DNA foci) and primarily ecDNA EGFR (>6 EGFR DNA foci). Statistical data relevant for this figure are in Table S5F. ns, not significant.

## References

Adelman, K., and Martin, B.J.E. (2021). ecDNA party bus: Bringing the enhancer to you. Mol. Cell 81, 1866–1867.

Boyle, S., Gilchrist, S., Bridger, J.M., Mahy, N.L., Ellis, J.A., and Bickmore, W.A. (2001). The spatial organization of human chromosomes within the nuclei of normal and emerin-mutant cells. Hum. Mol. Genet. 10, 211–219.

Brennan, C.W., Verhaak, R.G.W., McKenna, A., Campos, B., Noushmehr, H., Salama, S.R., Zheng, S., Chakravarty, D., Sanborn, J.Z., Berman, S.H., et al. (2013). The somatic genomic landscape of glioblastoma. Cell 155, 462–477.

Bulstrode, H., Johnstone, E., Marques-Torrejon, M.A., Ferguson, K.M., Bressan, R.B., Blin, C., Grant, V., Gogolok, S., Gangoso, E., Gagrica, S., et al. (2017). Elevated FOXG1 and SOX2 in glioblastoma enforces neural stem cell identity through transcriptional control of cell cycle and epigenetic regulators. Genes Dev. 31, 757–773.

Carvalho, C., Pereira, H.M., Ferreira, J., Pina, C., Mendonça, D., Rosa, A.C., and Carmo-Fonseca, M. (2001). Chromosomal G-dark bands determine the spatial organization of centromeric heterochromatin in the nucleus. Mol. Biol. Cell 12, 3563–3572.

Cho, W.-K., Spille, J.-H., Hecht, M., Lee, C., Li, C., Grube, V., and Cisse, I.I. (2018). Mediator and RNA polymerase II clusters associate in transcription-dependent condensates. Science 361, 412–415.

Chong, S., Dugast-Darzacq, C., Liu, Z., Dong, P., Dailey, G.M., Cattoglio, C., Heckert, A., Banala, S., Lavis, L., Darzacq, X., et al. (2018). Imaging dynamic and selective low-complexity domain interactions that control gene transcription. Science 361.

Cramer, P., Bushnell, D.A., Fu, J., Gnatt, A.L., Maier-Davis, B., Thompson, N.E., Burgess, R.R., Edwards, A.M., David, P.R., and Kornberg, R.D. (2000). Architecture of RNA polymerase II and implications for the transcription mechanism. Science 288, 640–649.

Croft, J.A., Bridger, J.M., Boyle, S., Perry, P., Teague, P., and Bickmore, W.A. (1999). Differences in the localization and morphology of chromosomes in the human nucleus. J. Cell Biol. 145, 1119–1131.

Deshpande, V., Luebeck, J., Nguyen, N.-P.D., Bakhtiari, M., Turner, K.M., Schwab, R., Carter, H., Mischel, P.S., and Bafna, V. (2019). Exploring the landscape of focal amplifications in cancer using AmpliconArchitect. Nat. Commun. 10, 392.

Dewari, P.S., Southgate, B., Mccarten, K., Monogarov, G., O’Duibhir, E., Quinn, N., Tyrer, A., Leitner, M.-C., Plumb, C., Kalantzaki, M., et al. (2018). An efficient and scalable pipeline for epitope tagging in mammalian stem cells using Cas9 ribonucleoprotein. Elife 7.

Dobin, A., Davis, C.A., Schlesinger, F., Drenkow, J., Zaleski, C., Jha, S., Batut, P., Chaisson, M., and Gingeras, T.R. (2013). STAR: ultrafast universal RNA-seq aligner. Bioinformatics 29, 15–21.

Fan, Y., Mao, R., Lv, H., Xu, J., Yan, L., Liu, Y., Shi, M., Ji, G., Yu, Y., Bai, J., et al. (2011). Frequency of double minute chromosomes and combined cytogenetic abnormalities and their characteristics. J. Appl. Genet. 52, 53–59.

Gangoso, E., Southgate, B., Bradley, L., Rus, S., Galvez-Cancino, F., McGivern, N., Güç, E., Kapourani, C.-A., Byron, A., Ferguson, K.M., et al. (2021). Glioblastomas acquire myeloid-affiliated transcriptional programs via epigenetic immunoediting to elicit immune evasion. Cell 184, 2454–2470.e26.

Heinz, S., Benner, C., Spann, N., Bertolino, E., Lin, Y.C., Laslo, P., Cheng, J.X., Murre, C., Singh, H., and Glass, C.K. (2010). Simple combinations of lineage-determining transcription factors prime cis-regulatory elements required for macrophage and B cell identities. Mol. Cell 38, 576–589.

Hung, K.L., Yost, K.E., Xie, L., Shi, Q., Helmsauer, K., Luebeck, J., Schöpflin, R., Lange, J.T., Chamorro González, R., Weiser, N.E., et al. (2021). ecDNA hubs drive cooperative intermolecular oncogene expression. Nature 1–6.

Imada, T., Shimi, T., Kaiho, A., Saeki, Y., and Kimura, H. (2021). RNA polymerase II condensate formation and association with Cajal and histone locus bodies in living human cells. Genes Cells 26, 298–312.

Inda, M.-M., Bonavia, R., Mukasa, A., Narita, Y., Sah, D.W.Y., Vandenberg, S., Brennan, C., Johns, T.G., Bachoo, R., Hadwiger, P., et al. (2010). Tumor heterogeneity is an active process maintained by a mutant EGFR-induced cytokine circuit in glioblastoma. Genes Dev. 24, 1731–1745.

Jubb, A., and Boyle, S. (2020). Visualizing Genome Reorganization Using 3D DNA FISH. In In Situ Hybridization Protocols, B.S. Nielsen, and J. Jones, eds. (New York, NY: Springer US), pp. 85–95.

Kalhor, R., Tjong, H., Jayathilaka, N., Alber, F., and Chen, L. (2011). Genome architectures revealed by tethered chromosome conformation capture and population-based modeling. Nat. Biotechnol. 30, 90–98.

Kent, W.J., Sugnet, C.W., Furey, T.S., Roskin, K.M., Pringle, T.H., Zahler, A.M., and Haussler, D. (2002). The human genome browser at UCSC. Genome Res. 12, 996–1006.

Kim, H., Nguyen, N., Turner, K., Wu, S., Liu, J., Deshpande, V., Namburi, S., Chang, H.Y., Beck, C., Mischel, P., et al. (2019). Frequent extrachromosomal oncogene amplification drives aggressive tumors.

Kim, H., Nguyen, N.-P., Turner, K., Wu, S., Gujar, A.D., Luebeck, J., Liu, J., Deshpande, V., Rajkumar, U., Namburi, S., et al. (2020). Extrachromosomal DNA is associated with oncogene amplification and poor outcome across multiple cancers. Nat. Genet.

Koga, T., Li, B., Figueroa, J.M., Ren, B., Chen, C.C., Carter, B.S., and Furnari, F.B. (2018). Mapping of genomic EGFRvIII deletions in glioblastoma: insight into rearrangement mechanisms and biomarker development. Neuro. Oncol. 20, 1310–1320.

Lange, J.T., Chen, C.Y., Pichugin, Y., Xie, L., Tang, J., Hung, K.L., Yost, K.E., Shi, Q., Erb, M.L., Rajkumar, U., et al. (2021). Principles of ecDNA random inheritance drive rapid genome change and therapy resistance in human cancers.

Li, H., Handsaker, B., Wysoker, A., Fennell, T., Ruan, J., Homer, N., Marth, G., Abecasis, G., Durbin, R., and 1000 Genome Project Data Processing Subgroup (2009). The Sequence Alignment/Map format and SAMtools. Bioinformatics 25, 2078–2079.

Morton, A.R., Dogan-Artun, N., Faber, Z.J., MacLeod, G., Bartels, C.F., Piazza, M.S., Allan, K.C., Mack, S.C., Wang, X., Gimple, R.C., et al. (2019). Functional Enhancers Shape Extrachromosomal Oncogene Amplifications. Cell 179, 1330–1341.e13.

Nathanson, D.A., Gini, B., Mottahedeh, J., Visnyei, K., Koga, T., Gomez, G., Eskin, A., Hwang, K., Wang, J., Masui, K., et al. (2014). Targeted therapy resistance mediated by dynamic regulation of extrachromosomal mutant EGFR DNA. Science 343, 72–76.

Pollard, S.M., Yoshikawa, K., Clarke, I.D., Danovi, D., Stricker, S., Russell, R., Bayani, J., Head, R., Lee, M., Bernstein, M., et al. (2009). Glioma stem cell lines expanded in adherent culture have tumor-specific phenotypes and are suitable for chemical and genetic screens. Cell Stem Cell 4, 568–580.

Quinlan, A.R., and Hall, I.M. (2010). BEDTools: a flexible suite of utilities for comparing genomic features. Bioinformatics 26, 841–842.

Rai, A.K., Chen, J.-X., Selbach, M., and Pelkmans, L. (2018). Kinase-controlled phase transition of membraneless organelles in mitosis. Nature 559, 211–216.

Ramírez, F., Ryan, D.P., Grüning, B., Bhardwaj, V., Kilpert, F., Richter, A.S., Heyne, S., Dündar, F., and Manke, T. (2016). deepTools2: a next generation web server for deep-sequencing data analysis. Nucleic Acids Res. 44, W160–W165.

Richards, L.M., Whitley, O.K.N., MacLeod, G., Cavalli, F.M.G., Coutinho, F.J., Jaramillo, J.E., Svergun, N., Riverin, M., Croucher, D.C., Kushida, M., et al. (2021). Gradient of Developmental and Injury Response transcriptional states defines functional vulnerabilities underpinning glioblastoma heterogeneity. Nature Cancer 2, 157–173.

Sabari, B.R., Dall’Agnese, A., Boija, A., Klein, I.A., Coffey, E.L., Shrinivas, K., Abraham, B.J., Hannett, N.M., Zamudio, A.V., Manteiga, J.C., et al. (2018). Coactivator condensation at super-enhancers links phase separation and gene control. Science 361.

Strom, A.R., and Brangwynne, C.P. (2019). The liquid nucleome - phase transitions in the nucleus at a glance. J. Cell Sci. 132.

Suvà, M.L., Rheinbay, E., Gillespie, S.M., Patel, A.P., Wakimoto, H., Rabkin, S.D., Riggi, N., Chi, A.S., Cahill, D.P., Nahed, B.V., et al. (2014). Reconstructing and reprogramming the tumor-propagating potential of glioblastoma stem-like cells. Cell 157, 580–594.

Turner, K.M., Deshpande, V., Beyter, D., Koga, T., Rusert, J., Lee, C., Li, B., Arden, K., Ren, B., Nathanson, D.A., et al. (2017). Extrachromosomal oncogene amplification drives tumor evolution and genetic heterogeneity. Nature 543, 122–125.

Veatch, S.L., Machta, B.B., Shelby, S.A., Chiang, E.N., Holowka, D.A., and Baird, B.A. (2012). Correlation functions quantify super-resolution images and estimate apparent clustering due to over-counting. PLoS One 7, e31457.

Verhaak, R.G.W., Hoadley, K.A., Purdom, E., Wang, V., Qi, Y., Wilkerson, M.D., Miller, C.R., Ding, L., Golub, T., Mesirov, J.P., et al. (2010). Integrated genomic analysis identifies clinically relevant subtypes of glioblastoma characterized by abnormalities in PDGFRA, IDH1, EGFR, and NF1. Cancer Cell 17, 98–110.

Verhaak, R.G.W., Bafna, V., and Mischel, P.S. (2019). Extrachromosomal oncogene amplification in tumour pathogenesis and evolution. Nat. Rev. Cancer 19, 283–288.

Vicario, R., Peg, V., Morancho, B., Zacarias-Fluck, M., Zhang, J., Martínez-Barriocanal, Á., Navarro Jiménez, A., Aura, C., Burgues, O., Lluch, A., et al. (2015). Patterns of HER2 Gene Amplification and Response to Anti-HER2 Therapies. PLoS One 10, e0129876.

Wang, L.-B., Karpova, A., Gritsenko, M.A., Kyle, J.E., Cao, S., Li, Y., Rykunov, D., Colaprico, A., Rothstein, J.H., Hong, R., et al. (2021). Proteogenomic and metabolomic characterization of human glioblastoma. Cancer Cell 39, 509–528.e20.

Wu, S., Turner, K.M., Nguyen, N., Raviram, R., Erb, M., Santini, J., Luebeck, J., Rajkumar, U., Diao, Y., Li, B., et al. (2019). Circular ecDNA promotes accessible chromatin and high oncogene expression. Nature 575, 699–703.

Yi, E., Gujar, A.D., Guthrie, M., Kim, H., Zhao, D., Johnson, K.C., Amin, S.B., Costa, M.L., Yu, Q., Das, S., et al. (2021). Live-cell imaging shows uneven segregation of extrachromosomal DNA elements and transcriptionally active extrachromosomal DNA hubs in cancer. Cancer Discov.

Zhu, Y., Gujar, A.D., Wong, C.-H., Tjong, H., Ngan, C.Y., Gong, L., Chen, Y.-A., Kim, H., Liu, J., Li, M., et al. (2021). Oncogenic extrachromosomal DNA functions as mobile enhancers to globally amplify chromosomal transcription. Cancer Cell 39, 694–707.e7.

